# Live-cell imaging and physical modeling reveal control of chromosome folding dynamics by cohesin and CTCF

**DOI:** 10.1101/2022.03.03.482826

**Authors:** Pia Mach, Pavel I. Kos, Yinxiu Zhan, Julie Cramard, Simon Gaudin, Jana Tünnermann, Edoardo Marchi, Jan Eglinger, Jessica Zuin, Mariya Kryzhanovska, Sebastien Smallwood, Laurent Gelman, Gregory Roth, Elphège P. Nora, Guido Tiana, Luca Giorgetti

**Author notes:** equal contribution (Pavel I. Kos, Yinxiu Zhan).

## Abstract

Physical proximity between genomic sequences in mammalian chromosomes controls key biological processes such as transcriptional regulation and DNA repair. Yet it is currently unknown if chromosomal contacts are rare and stable or instead frequent and dynamic, and how they depend on the loop extrusion activity of cohesin or barriers such as CTCF. By imaging chromosomal locations at high spatial and temporal resolution over several hours in living cells, we show that sequences within topological associating domains (TADs) frequently come into physical proximity during the course of a cell cycle and remain close to each other only for a few minutes. Such contacts become nonetheless substantially longer and more frequent in the presence of convergent CTCF sites, resulting in a suppression of variability in chromosome folding in single cells across time. Supported by physical models of chromosome dynamics, our data additionally suggests that individual CTCF-anchored loops last around 10 minutes. The estimates of chromosomal contact dynamics in our study provide a novel quantitative framework to link chromosome structure to function and show that cohesin and CTCF stabilize otherwise highly dynamic chromosome structures to facilitate selected subsets of chromosomal interactions.

## Introduction

In the nuclei of mammalian cells, the physical interactions between sequences separated by large genomic distances play an important role in fundamental processes such as DNA replication^1^, repair^2^ and transcriptional regulation by distal enhancers^3^. Chromosome conformation capture (3C) methods, which measure physical proximity between genomic sequences in fixed cells, have revealed that chromosomal contacts are organized into sub-megabase domains of preferential interactions known as topologically associating domains (TADs)^4, 5^ whose boundaries are able to functionally insulate regulatory sequences^3^ . TADs mainly arise from nested interactions between convergently oriented binding sites of the DNA-binding protein CTCF. This is thought to occur as chromatin-bound CTCF arrests the loop-extrusion activity of the cohesin complex^6–8^. Depletion of CTCF or key components of the cohesin complex, such as RAD21, results in loss of TADs^9–11^ . In contrast, degradation of the cohesin release factor WAPL results in the stabilization of cohesin on DNA and in an increase of interactions between CTCF sites^10, 12^.

Determining the temporal dynamics of chromosome structure and its relationship with CTCF and cohesin is fundamental for achieving a quantitative understanding of nuclear processes and notably long-range transcriptional regulation^3, 13–15^. Single-cell analyses of chromosome structure in fixed cells^16–18^ as well as *in vitro*^19^ and live-cell^20^ measurements of CTCF and cohesin dynamics have suggested that CTCF loops and TADs are dynamic structures^14^. However we have little knowledge on whether and how loop extrusion contributes to chromosome dynamics in living cells, and impacts the rates and durations of encounters between genomic sequences, especially at the scale of TADs. It is unclear if in physiological conditions cohesin would increase encounter rates between genomic sequences *in vivo* by reeling them into loops, or rather prolong the duration of their encounters, or both. Recent analysis of a 500-kb CTCF loop connecting opposite TAD boundaries provided initial constraints on the duration and frequency of cohesin/CTCF-mediated interactions^21^. Yet the frequencies and durations of chromosomal interactions inside TADs and their relationship with the loop extrusion process have not been systematically investigated in living cells.

Here we use live-cell fluorescence microscopy to measure chromosome dynamics and its dependence on cohesin and CTCF in mouse embryonic stem cells (mESC). By combining two live-cell imaging strategies with polymer simulations, we reveal that loops extruded by cohesin constrain global chromosome motion, while also increasing the temporal frequencies and durations of physical encounters between sequences inside TADs. Convergent CTCF sites substantially stabilize contacts between sequences through cohesin-mediated CTCF-anchored loops that last around 5 to 15 minutes on average. Our results support the notion that chromosome structure within single TADs is highly dynamic during the span of a cell cycle. They also reveal how contact dynamics and the temporal variability in chromosome folding are modulated by cohesin and CTCF in single living cells and provide a quantitative framework for understanding the role of folding dynamics in fundamental biological processes.

### Cohesin substantially decreases global chromosome mobility in living cells independent of CTCF

To study how cohesin and CTCF influence the global dynamics of the chromatin fiber independent of the local chromatin state and structural differences, we first studied the dynamic properties of large numbers of random genomic locations in living cells. To this aim, we generated clonal mESC lines carrying multiple random integrations of an array of approximately 140 repeats of the bacterial Tet operator sequence (TetO) using piggyBac transposition^22^. These can be visualized upon binding of Tet repressor (TetR) fused to the red fluorescent protein tdTomato^23^. To compare the motion of genomic locations that either block or allow the loop extrusion activity of cohesin, the TetO array was adjacent to a cassette carrying three CTCF motifs (3xCTCF) that could be removed by Cre-assisted recombination **(Fig. 1A)**. Motifs were selected based on high CTCF enrichment in ChIP-seq and each was confirmed to be bound by CTCF in nearly 100% of alleles at any time in mESCs using dual-enzyme single-molecule footprinting^24^ (R. Grand and D. Schubeler, personal communication), thus providing a close experimental representative of an “impermeable” loop extrusion barrier.

**Figure 1:**
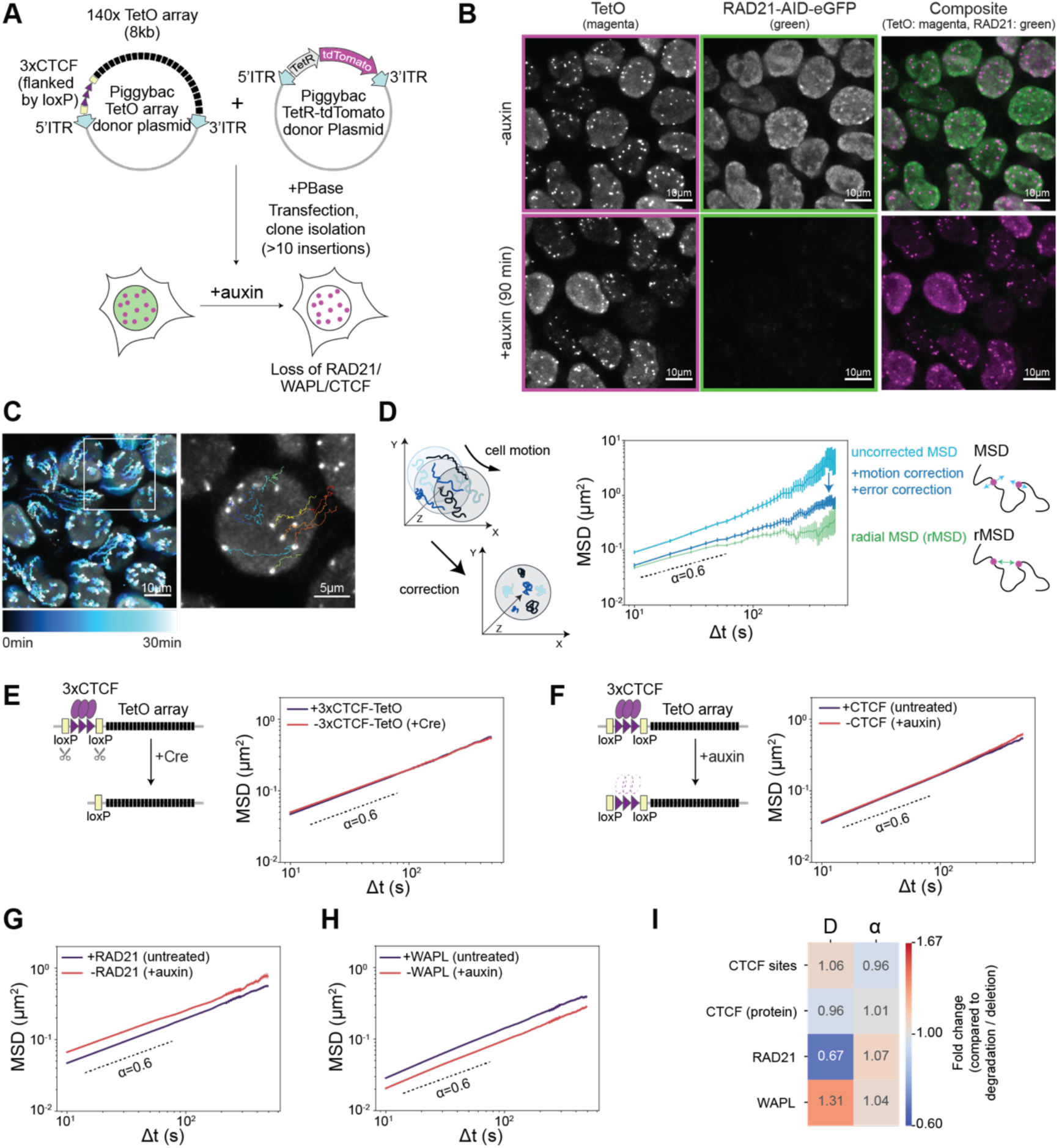
Cohesin decelerates chromosome dynamics in living cells. A. Generation of clonal mESC lines containing random integrations of Tet operator (TetO) arrays flanked by 3xCTCF motifs and carrying TetR-tdTomato for visualization of TetO arrays. All constructs were integrated using piggyBac transposition in mESCs allowing auxin-inducible degradation of either GFP-tagged RAD21, WAPL or CTCF. ITR: inverted terminal repeats. B. Representative images of RAD21-AID-eGFP containing 3xCTCF-TetO cells imaged either before (-auxin) or after 90 min (+auxin) of treatment (exposure time (eGFP) = 50 ms, exposure time (tdTomato) = 50ms, deconvolved, max. intensity projection, bicubic interpolation). C. Left: 2D projection of an images series of TetR-tdTomato signal over 30 min (dt=10 s, color-coded for intensity changes over time. Zoom-in: Overlay of TetR-tdTomato signal with reconstructed trajectories of individual TetO arrays. D. Left: Cell motion is approximated at every time frame as the average roto-translational motion of the ensemble of trajectories within the same nucleus. Right: Mean Square Displacement (MSD) averaged over individual trajectories within one nucleus (MSD, mean +/- standard error) before (cyan) and after (dark blue) cell motion and localisation error correction. Green: radial MSD of pairs of operator arrays within the same nucleus (rMSD, mean +/- standard error, green). E. Left: Cre-mediated excision of the 3xCTCF cassette preserves genomic positions of the arrays. Right: MSD (mean +/- standard error) in mESC lines in presence (blue line: 310 cells, 13,537 trajectories) or absence (red line: 271 cells, 11,082 trajectories) of 3xCTCF next to TetO arrays. Three replicates per clonal cell line and three clonal lines per condition were analyzed and merged in this graph and all following MSD graphs. P-values (two-sided Student t-test) for all panels shown in **Suppl. Fig. S2E**. F. Left: Acute depletion of CTCF in CTCF-AID-eGFP cells induced by treatment with auxin for 6 h. Right: Same as in E right panel, but in mESC lines with 3xCTCF-TetO arrays before (blue line: 323 cells, 9,829 trajectories analyzed) and after CTCF degradation (red line: 365 cells, 12,495 trajectories analyzed). G. MSD (mean +/- standard error) of 3xCTCF-TetO insertions before (blue line: 310 cells, 13,537 trajectories) or after RAD21 degradation upon 90 minutes treatment with auxin (red line: 240 cells, 8,788 trajectories). H. MSD (mean +/- standard error) of 3xCTCF-TetO before (blue line: 336 cells, 6,687 trajectories) or after WAPL degradation upon 24 hours treatment with auxin (red line: 350 cells, 6,717 trajectories analyzed). I. Heatmap representing fold changes of generalized diffusion coefficients (D) and scaling exponents (*α*) in untreated cells compared to conditions when the corresponding factor is degraded (CTCF, RAD21, WAPL) or the CTCF motifs (3xCTCF) are deleted.

3xCTCF-TetO sequences were introduced in mESC lines that stably expressed *Os*Tir1 and where the endogenous *Rad21*, *Wapl* or *Ctcf* genes were targeted with an auxin-inducible degron (AID) peptide fused to eGFP^7, 9, 25^. This resulted in several mESC clones (three per degron condition), each with different sets of genomic insertions of the 3xCTCF-TetO cassette, where over 95% of any of the AID-tagged proteins could rapidly be depleted from the nucleus upon addition of auxin (**Fig. 1B** and **Suppl. Fig. S1A**). This allowed us to study chromosome dynamics after the acute depletion of factors affecting cohesin-mediated chromosome structure (**Suppl. Fig. S1B**) at previously reported timepoints (90 minutes for RAD21^11^, 6 hours for CTCF^9^ and 24 hours for WAPL^25^) that minimize secondary effects such as defects in cell-cycle progression(**Suppl. Fig. S1C)**,).

Mapping of the TetO insertion sites in the clonal lines revealed that there were 10 to 20 insertions per cell line, with on average 1 or 2 heterozygous insertions per chromosome without any strong bias towards active or inactive chromatin **(Suppl. Fig. S1D,E)**. Insertion sites were on average at a 10-kb distance from the nearest endogenous CTCF binding sites (**Suppl. Fig. S1F**). 4C-seq confirmed that insertion of the 3xCTCF-TetO cassette often led to the formation of ectopic interactions with endogenous CTCF sites, which were lost upon removal of the 3xCTCF sites or depletion of RAD21 (**Suppl. Fig. S1G,H**).

To measure the dynamic properties of 3xCTCF-TetO insertions, we acquired 3D movies (one z-stack of 10 µm every 10 seconds for 30 minutes) using highly inclined and laminated optical sheet (HILO) microscopy^26^ **(Fig. 1B, Suppl. Video S1)**. This resulted in approx. 270 cells/condition with over 8,000 trajectories from three clonal lines imaged with 3-4 biological replicates for each condition. Detection and localisation of TetO arrays as sub-diffraction fluorescent signals^27^ enabled reconstruction of the trajectories of individual genomic insertions (**Fig. 1C, Methods**). We then studied their mean squared displacement (MSD) as a function of time after correcting each trajectory for the confounding effect of cell movement, which we inferred from the collective displacement of all the insertions in each nucleus (**Fig. 1D, Suppl. Fig. S2A,** see **Methods)**. Independent of the degron background, in untreated cells genomic locations underwent on average a subdiffusive motion whose anomalous exponent (approx. 0.6) and generalized diffusion coefficients D (approx. 1.2×10^-2^ µm^2^s^-^*α*) were in line with previous studies of specific genomic loci^28, 29^ **(Fig. 1D, Suppl. Fig. S2B)**. The MSD of radial distances (radial MSD, rMSD) between insertions within the same nuclei showed the same scaling although the statistics were less robust for long time intervals due to the shorter trajectories that could be built based on pairwise distances **(Fig. 1D)**. Interestingly, removal of the 3xCTCF sites **(Suppl. Fig. S1G)** or degradation of CTCF (6 hours auxin treatment in CTCF-AID cells) did not have a significant impact on MSD averaged over all genomic locations nor on its distribution across trajectories and cells (**Fig. 1E,F**, results for single clones in **Suppl. Fig. S2C,** p-values in **Suppl. Fig. S2E**).

By contrast, acute depletion of RAD21 (90 min treatment with auxin) led to a significant increase in mobility of insertions both in the presence **(Fig. 1G)** and absence of 3xCTCF sites (**Suppl. Fig. S2D**), with only a very minor impact on anomalous exponents **(Suppl. Fig. S2B,E,** p-values in **Suppl. Fig. S2E)**. In the presence of wild-type levels of RAD21, generalized diffusion coefficients were on average approximately 30% lower than in depleted cells, in which RAD21 levels were low enough to prevent formation of cohesin-mediated structures (**cf. Suppl. Fig. S1B**). This outcome was consistent across three clonal cell lines with different TetO insertion sites and the small differences in the magnitude of the effect were likely due to location-dependent effects (**Suppl. Fig. S2F**). Importantly, the effect was specific for RAD21 degradation as we did not observe any changes in MSD behavior in control cell lines expressing *Os*Tir1 but no AID-tag **(Suppl. Fig. S2G)**. In addition, depletion of the cohesin release factor WAPL, which results in higher levels of DNA-bound cohesin^25^, caused a substantial decrease in chromosome mobility (24 hours auxin treatment of WAPL-AID clones) (**Fig. 1H, Suppl. Fig. S2E)**. Together these results thus indicate that increasing levels of DNA-bound cohesin decelerate chromosome motion, with only very minor effects (if any) mediated by the presence of even strong CTCF motifs (**Fig. 1I**).

### Loop extrusion explains the decelerating effect of cohesin on chromosome dynamics

We next used polymer simulations to determine if the process of loop extrusion alone could explain the observed global deceleration of chromosome motion in the presence of cohesin and the minimal effects from CTCF sites. We simulated the dynamics of a polymer chain with excluded volume, with or without loop extrusion and extrusion barriers whose linear arrangement and orientation was sampled from endogenous CTCF sites (**Fig. 2A, Suppl. Fig. S3A**). To mimic the random insertion of 3xCTCF sites, we also simulated the same polymers with additional randomly inserted loop extrusion barriers (zoom-in in **Fig. 2A**). To emphasize their potential effects on chromosome dynamics, all barriers in the simulations were impermeable to loop extrusion. Every monomer represented 8 kb of chromatin, corresponding to the genomic size of the TetO array and simulation steps were converted into real time units by matching the time needed for a monomer and TetO array to move by their own sizes (see **Methods**). We sampled an extremely large range of extruder residence times and loading rates (4 orders of magnitude each) centered around a residence time of ∼30 min and extruder densities of ∼20 per Mb (in line with previous measurements^30, 31^), and using two extrusion speeds corresponding to *in vivo* and *in vitro* estimates (∼0.1 kb/sec and ∼1 kb/sec, respectively)^19, 30^ (**Fig. 2B, Suppl. Fig. S3B**).

**Figure 2:**
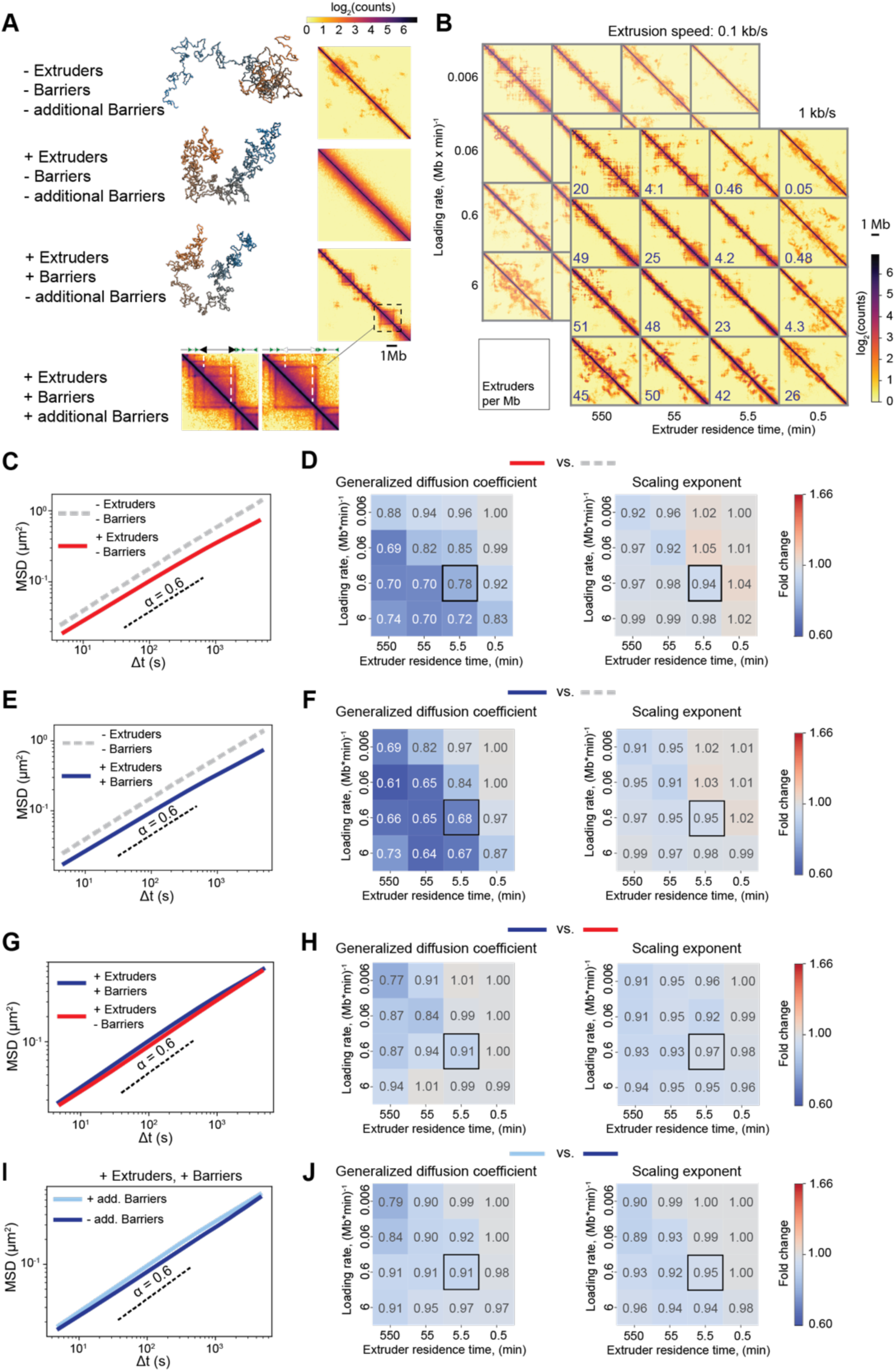
Simulations predict that loop extrusion generally decelerates polymer motion. A. Representative snapshots of conformations and simulated contact maps for a polymer model with excluded volume and increasingly complex models with loop extruders, extrusion barriers sampled from CTCF motifs within eight Mb on chromosome 15 (Chr15:7-16Mb), and additional randomly distributed extrusion barriers. For the system with additional barriers, contact map is presented aside with zoom-in of the contact map of the system without additional barriers to highlight the differences. B. Simulated contact maps (with loop extrusion and extrusion barriers) for polymers with two extrusion speeds (1 kb/s and 0.1 kb/s) and different combinations of extruder loading rates and residence times. The resulting linear densities of extruders (number per Mb) are shown in the bottom left corner of each contact map. C. Effect of extruders. MSDs of polymers with (red line) or without (gray dashed line) loop extruders in the absence of extrusion barriers (loading rate 0.6 (Mb x min)^-1^ and residence time 5.5 min, corresponds to black frame in panel D). Dashed curve represents *α* = 0.6 as an eyeguide. D. Effect of extruders.Ratios of generalized diffusion coefficients and anomalous exponents between the two conditions shown in panel C. Black square: set of parameters whose corresponding MSDs are shown in panel C. E. MSDs of polymers with (blue line) or without (gray dashed line) both extruders and barriers. Same parameters as in panel C. F. Same as panel D for cases illustrated in panel E. G. MSDs of polymers with loop extruders in the presence (blue) or absence (red) of extrusion barriers. Same parameters as in panels C and E. H. Same as panels D and F but for cases illustrated in panel G. I. MSDs of polymers either with (light blue) or without (red) additional randomly inserted extrusion barriers. Same parameters as in panels C, E, G. J. Same as panels D, F and H but for cases illustrated in panel I.

In the absence of loop extrusion, the polymer underwent subdiffusive behavior with an anomalous exponent of approx. 0.6 (**Fig. 2C**), as expected from a simple polymer with excluded volume^32^ (see **Supplementary Material**) and compatible with our experimental results on random TetO insertions (**Fig. 1E**). Strikingly, in line with the experimental effects of RAD21 (**Fig. 1G**), introduction of loop extrusion led to lower generalized diffusion coefficients and very minor effects on anomalous exponents, independent of extruder loading rate and residence time (**Fig. 2C,D**), extrusion speed (**Suppl. Fig. S3E**), or the presence of extrusion barriers (**Fig. 2E,F**). Interestingly, for extruder residence times of 5.5-11 minutes and unloading rates corresponding to extruder linear densities of ∼20 per Mb, the predicted decrease in generalized diffusion coefficients was in quantitative agreement with the experimentally observed value of approx. 30% (**Fig. 2D,F**; extruder densities as in **Fig. 2B**; cf. **Fig. 1G**). Also, consistent with WAPL depletion experiments (**Fig. 1H**), increasing extruder residence times systematically resulted in larger reductions in generalized diffusion coefficients (**Fig. 2D, F, Suppl. Fig S3E**).

Importantly, addition of barriers in the presence of loop extrusion led to substantially smaller changes in polymer dynamics compared to the effect of loop extrusion itself even when probed directly on the barriers (**Fig. 2G,H, Suppl. Fig. S3C-E**), in agreement with our experimental finding that CTCF degradation had no strong effect on the MSD of TetO insertions (**Fig. 1E, F**,**Suppl. Fig. S2C,E**). Similarly, insertion of additional barriers had little impact on MSD (**Fig. 2I-J**), thus recapitulating the experimental negligible effect of removal of 3xCTCF sites (**Fig. 1E**). Polymer simulations thus strongly support the notion that the observed decrease in chromosome mobility and lack of substantial effects from strong CTCF motifs is a general, macroscopic manifestation of the loop extrusion process in living cells.

### Both cohesin and CTCF constrain the reciprocal dynamics of sequences within the same TAD

The finding that CTCF does not impact global chromosome mobility is at odds with 3C-based findings that sites bound by CTCF form site-specific interactions with increased contact probabilities. We therefore asked how cohesin and CTCF impact the reciprocal motion of two genomic sequences located on the same DNA molecule. To this aim we simulated the dynamics of a polymer carrying two impermeable convergent extrusion barriers mimicking strong CTCF motifs separated by approximately 150 kb (**Fig. 3A**). This is comparable to the median distance between convergent CTCF sites within TADs genome-wide in mESC (141 kb, see **Methods**) and also to the estimated average separation between enhancers and promoters in human cells (∼160 kb)^33^. Simulations performed with extrusion parameters recapitulating the dynamic effect of RAD21 depletion (black square in **Fig. 2F**) predicted that rMSDs should be lowest in the presence of loop extrusion and extrusion barriers (**Fig. 3B**) due to the formation of transient loops anchored by the barriers (**Fig. 3C**). Similar to the behavior observed for MSDs (**Fig. 2**), rMSDs should increase upon removal of extrusion barriers and become maximal when loop extrusion is also removed (**Fig. 3B**).

**Figure 3:**
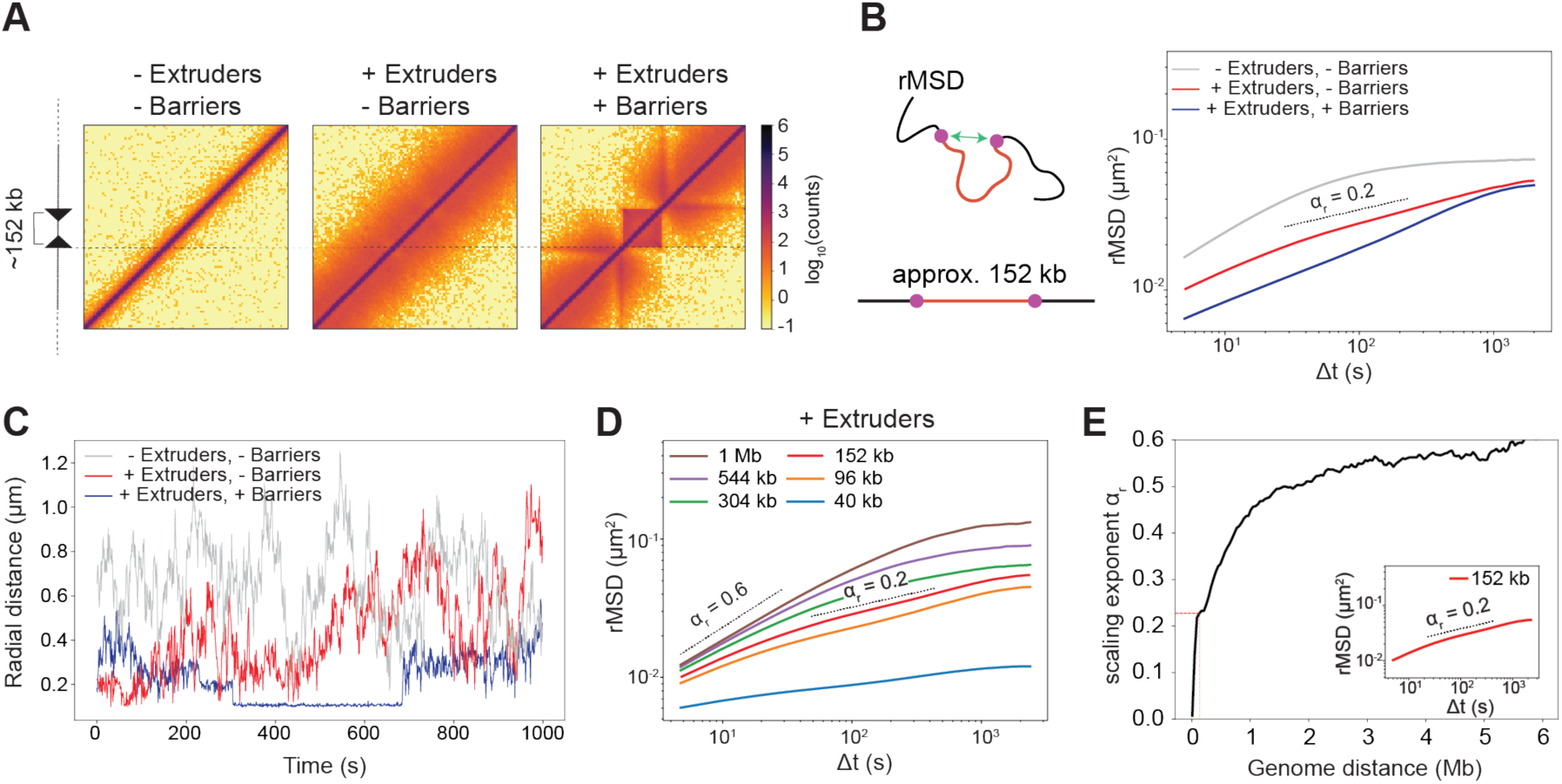
Polymer simulations of the reciprocal dynamics of two sequences in *cis*. A. Simulated contact maps of a region spanning the equivalent of 800 kb for a polymer chain without loop extrusion, with loop extruders, and with convergent extrusion barriers separated by the equivalent of 152 kb. B. Radial MSD (rMSD) of the two monomers separated by the equivalent of 152 kb in the three conditions from panel A. Dashed line is an exponent 0.2 serving as an eyeguide (*α*r indicates the slope of rMSDs). Loop extrusion parameters as in Fig. 2C. C. Representative examples of distances between the two monomers in simulations with or without loop extrusion and extrusion barriers. The flat stretch in the trajectory with extrusion and barriers corresponds to a loop anchored by the two barriers. D. rMSD of distances between multiple pairs of monomers separated by distances equivalent to 40 Kb - 1 Mb for a polymer with loop extrusion but no barriers. E. Slopes of rMSD curves for two loci separated by varying linear distances, estimated from linear fitting between 5 and 60 seconds. Inset: detail of rMSD and fit for monomers separated by 152 kb.

Importantly, simulations also predicted that scaling exponents of rMSD curves should be considerably smaller (around 0.2) than those we previously observed for TetO arrays separated by several Mb or located on different chromosomes (around 0.6, **Fig. 1D**). This is because correlations in the motion of two monomers are much stronger when they are located nearby along the polymer than when they are located far away. Indeed, simulations predicted that the scaling exponent fitted from rMSD curves at short times should increase with increasing genomic distance and approach 0.6 for loci separated by several Mb (consistent with rMSDs of randomly inserted TetO arrays) (**Fig. 3D,E**) before saturating to stationary values at longer times. This holds true without loop extrusion (see theoretical analysis in **Suppl. Material** and simulations in **Suppl. Fig. S4A,B**).

To test these predictions, we turned to a live-cell imaging approach allowing us to measure the radial dynamics of two sequences located within the same TAD, in the presence and absence of cohesin and/or strong CTCF sites. Specifically, we engineered a mESCs line carrying targeted integrations of two orthogonal operator arrays (approx. 140x TetO and 120x LacO separated by 150 kb (**Fig. 4A**), which could be visualized upon binding of TetR-tdTomato and a weak DNA-binding variant of LacI fused to eGFP (LacI**-eGFP)^23^. To minimize confounding effects from additional regulatory sequences such as expressed genes or active enhancers, we targeted the arrays into a 560-kb “neutral” TAD on chromosome 15 which we also previously engineered to remove internal CTCF binding sites^3^ (**Fig. 4A**). The two operator arrays were directly adjacent to excisable 3xCTCF site cassettes arranged in a convergent orientation (**Fig. 4A**). Cell lines were verified by Nanopore Cas9-targeted sequencing (nCATS)^34^ to contain a single copy of each targeting cassette (**Suppl. Fig. S4C**). We additionally targeted the endogenous *Rad21* locus with a C-terminal HaloTag-FKBP fusion allowing the inducible degradation of RAD21 upon treatment with dTAG- 13^35^ as confirmed by severely decreased protein levels (>95% after 2 hours of treatment, **Suppl. Fig. S4D**).

**Figure 4:**
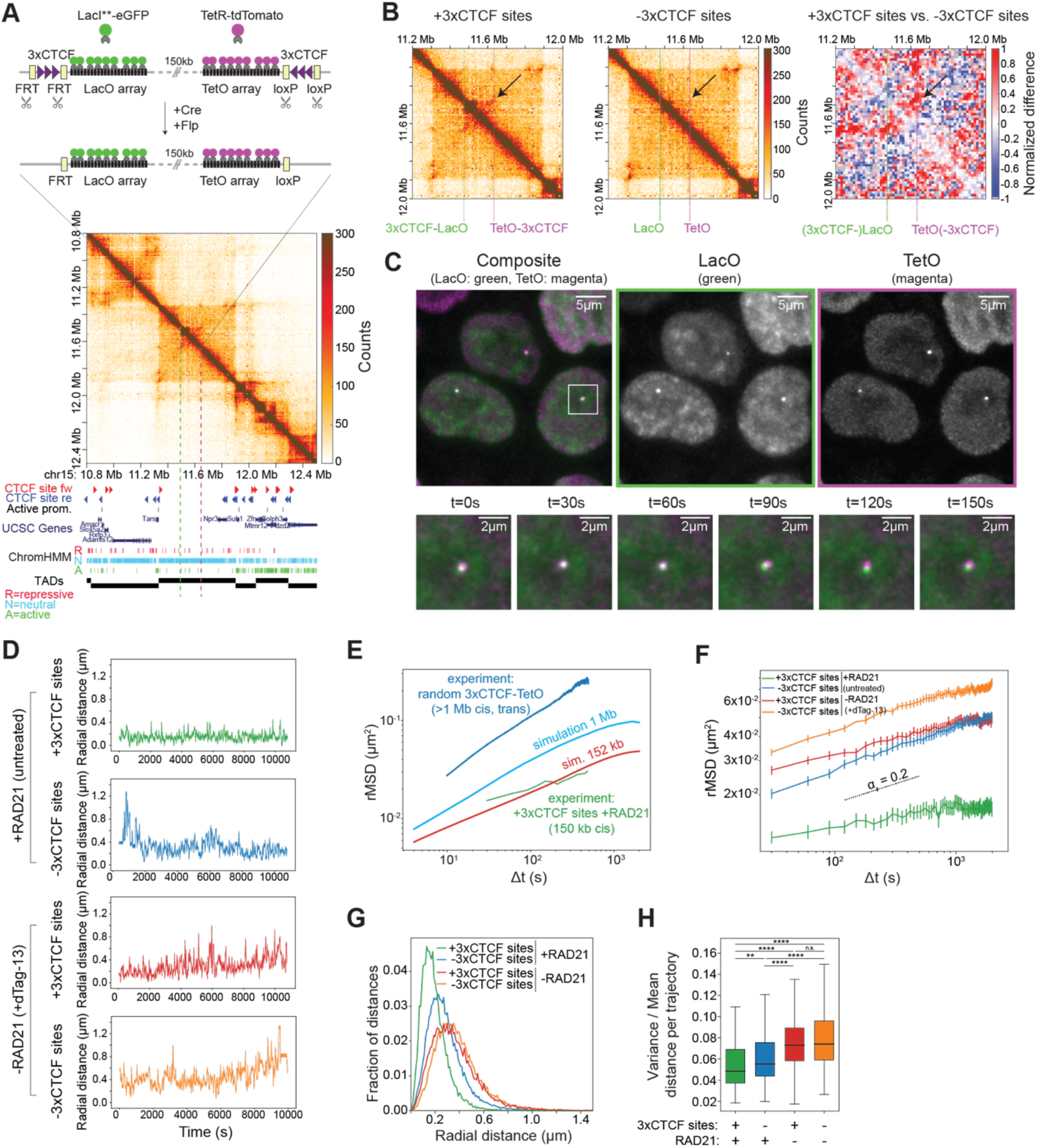
Live-cell imaging of two sequences within the same TAD, with or without cohesin and or CTCF sites. A. Top: insertion of TetO and LacO arrays separated by 150 kb within a “neutral” TAD on chromosome 15 in mESC. 3xCTCF sites flanking the arrays can be excised by Cre and Flp recombinases. Arrays were visualized by binding of Lac**-eGFP and TetR-tdTomato, respectively. Bottom: tiled Capture- C map (6.4 kb resolution) plotted with genomic datasets in mESC in a region of 2.6 Mb surrounding the engineered TAD. Capture-C was performed in cells where arrays were flanked by 3xCTCF sites. Dashed magenta and green lines: positions of LacO and TetO insertions. B. Capture-C maps in mESC lines with (left) or without (middle) 3xCTCF sites flanking the TetO and LacO arrays, and differential map (right, +3xCTCF vs. -3xCTCF, see Methods) highlighting that convergent 3xCTCF sites lead to the formation of interactions between the two insertion locations (arrows). C. Top: Representative fluorescence microscopy images of mESCs with 3xCTCF-LacO and TetO- 3xCTCF insertions. Bottom: Zoomed-in view showing a time-series overlay of LacI**-eGFP and TetR-tdTomato signals (exposure time 50 ms, deconvolved, max. intensity projection, bicubic interpolation). D. Representative trajectories of TetO-LacO radial distances in mESC lines with or without convergent 3xCTCF sites, either before or after treatment with 500 nM dTag-13 for 2 hours to induce degradation of RAD21 (dt=30s). E. rMSD of 3xCTCF-TetO random integrations (dark blue, see Fig. 1E) and of targeted 3xCTCF-LacO and TetO-3xCTCF insertions on Chr15 (green) are compared to model predictions for pairs of loci containing extrusion barriers at a distance of 1Mb (light blue) and 152 kb (red). Note that random 3xCTCF-TetO insertions often occur on different chromosomes and thus have larger absolute rMSD than 1Mb simulations (but similar scaling). F. rMSD of TetO-LacO distances in cell lines in mESC lines with or without convergent 3xCTCF sites, either before or after treatment with 500 nM dTag-13 for 2 hours to induce degradation of RAD21 (dt=30s). rMSDs are plotted as mean +/- standard error over n=4-7 replicates for each condition. No. of cells analyzed: +CTCF sites/+RAD21: 152 (4 replicates), -CTCF sites/+RAD21: 214 (4 replicates), +CTCF sites/-RAD21: 248 (7 replicates), -CTCF sites/-RAD21: 277 (6 replicates). G. Distribution of TetO-LacO radial distances in the four experimental conditions (n=4-7 replicates per conditions, as in panel F). H. Distributions of variance over mean of single trajectories across the four experimental conditions (n=4-7 replicates per conditions, as in panel F). Boxes: lower and upper quartiles (Q1 and Q3, respectively). Whiskers: denote 1.5x the interquartile region (IQR) below Q1 and above Q3. P-values are calculated using Kolmogorov–Smirnov test, two-sided: n.s. - not significant, ** - p<0.01, **** - p<0.0001.

Capture-C with tiled oligonucleotides revealed that integration of the operator arrays themselves did not lead to detectable changes in chromosome structure (**Suppl. Fig. S4E**). Convergent 3xCTCF sites however led to the formation of a novel CTCF-mediated interaction within the TAD (2.6-fold increase in contact probability after correction of the confounding contribution of the wild-type non-targeted allele) (**Fig. 4B**), which was lost upon depletion of RAD21 along with all other CTCF-mediated interactions across the locus (**Suppl. Fig. S4F**).

We imaged cells for three hours every 30 seconds in 3D (**Fig. 4C, Suppl. Video S2**) either in the presence or absence of RAD21 and measured radial distances between the two operator arrays over time (**Fig. 4D,** n=4-7 biological replicates for each condition, on average 223 cells analyzed per condition, for details see **Suppl. Table S1** and **Methods**). Doublet signals corresponding to replicated alleles occurred in a very minor fraction (3%, see **Methods**) of trajectories, compatible with the late-replication profile of the “neutral” TAD and the cell-cycle distribution (**Suppl. Fig. S4G,H**). In these cases, only trajectories that were initially closest across channels were considered. We estimated our experimental uncertainty on radial distances after correction of chromatic aberrations (see **Methods**) to be approximately 130 nm by measuring pairwise distances in a control cell line where multiple TetO insertions were simultaneously bound by both TetR-tdTomato and TetR-eGFP (**Suppl. Fig. S4I-K**).

In agreement with model predictions for locations separated by 150 kb, rMSDs for the two operator arrays showed a low scaling behavior, with exponents close to 0.2 and thus much smaller than those observed with randomly inserted TetO arrays (**Fig. 4E,F**). Also in line with model predictions (**Fig. 3D**), the presence of RAD21 and 3xCTCF sites led to the most constrained radial mobility, whereas RAD21 degradation and deletion of CTCF sites resulted in the least constrained motion (**Fig. 4F**). These measurements thus verified the model prediction that genomic sequences located at short distance (150 kb) in *cis* experience stronger physical constraints than sequences located at larger genomic distances^21^ (**Fig. 1, Suppl. Fig. S4L**), and that loop extrusion provides constraints that are further reinforced by the presence of convergent CTCF sites.

Consistent with their more constrained rMSD behavior, we finally observed that radial distances between TetO and LacO signals were smallest in the presence of convergent CTCF sites and cohesin. In these conditions, distances between TetO and LacO arrays tended to remain close to the ∼130 nm experimental uncertainty on radial distances with only occasional fluctuations towards larger values in the course of the 3 hours of imaging (**Fig. 4D,G**). Removal of 3xCTCF sites led to an increase in radial distances and in their variability within single trajectories, which were further increased upon degradation of RAD21, irrespective of the presence or absence of CTCF sites (**Fig. 4D,G, Suppl. Video S3**). Thus constraints imposed by extruding cohesin and convergent CTCF sites reduce not only the average physical distances between sequences but also their variability in time (**Fig. 4H**).

### Chromosomal contacts are transient and stabilized by cohesin and CTCF

We next set off to quantify changes in distances over time and determine whether despite the experimental uncertainty on 3D distances (**Suppl. Fig. S4K**) we could observe transitions between two states: a “proximal” state with small radial distances (presumably including cohesin-mediated loops between convergent CTCF sites), and a generic “distal” state with larger spatial distances corresponding to other configurations of the chromatin fiber. To this aim we fitted a two-state hidden Markov model (HMM) on the ensemble of trajectories obtained in cells where both convergent 3xCTCF sites and RAD21 were present (**Fig. 5A, top left, Fig. 5B**). Interestingly, distances in the proximal state inferred by HMM largely overlapped with those detected on perfectly colocalizing signals in control experiments (149 vs. 130 nm on average, respectively) (**Fig. 5B right panel, cf. Suppl. Fig. S4K**). The proximal state thus corresponds to configurations of the chromatin fiber where the two arrays were in very close physical proximity, also including (but not restricted to) cohesin-mediated loops between CTCF sites. For simplicity we will refer to the proximal state interchangeably as ‘contact’, without implying a direct molecular interaction between the two DNA fibers. Radial distances in the distal state (288 nm on average) instead were similar to those measured in cells where both CTCF sites had been removed (291 nm) (**Suppl. Fig. S5A,B)**. Thus the distal state largely overlapped with chromosome conformations where specific cohesin-mediated CTCF loops were lost.

**Figure 5:**
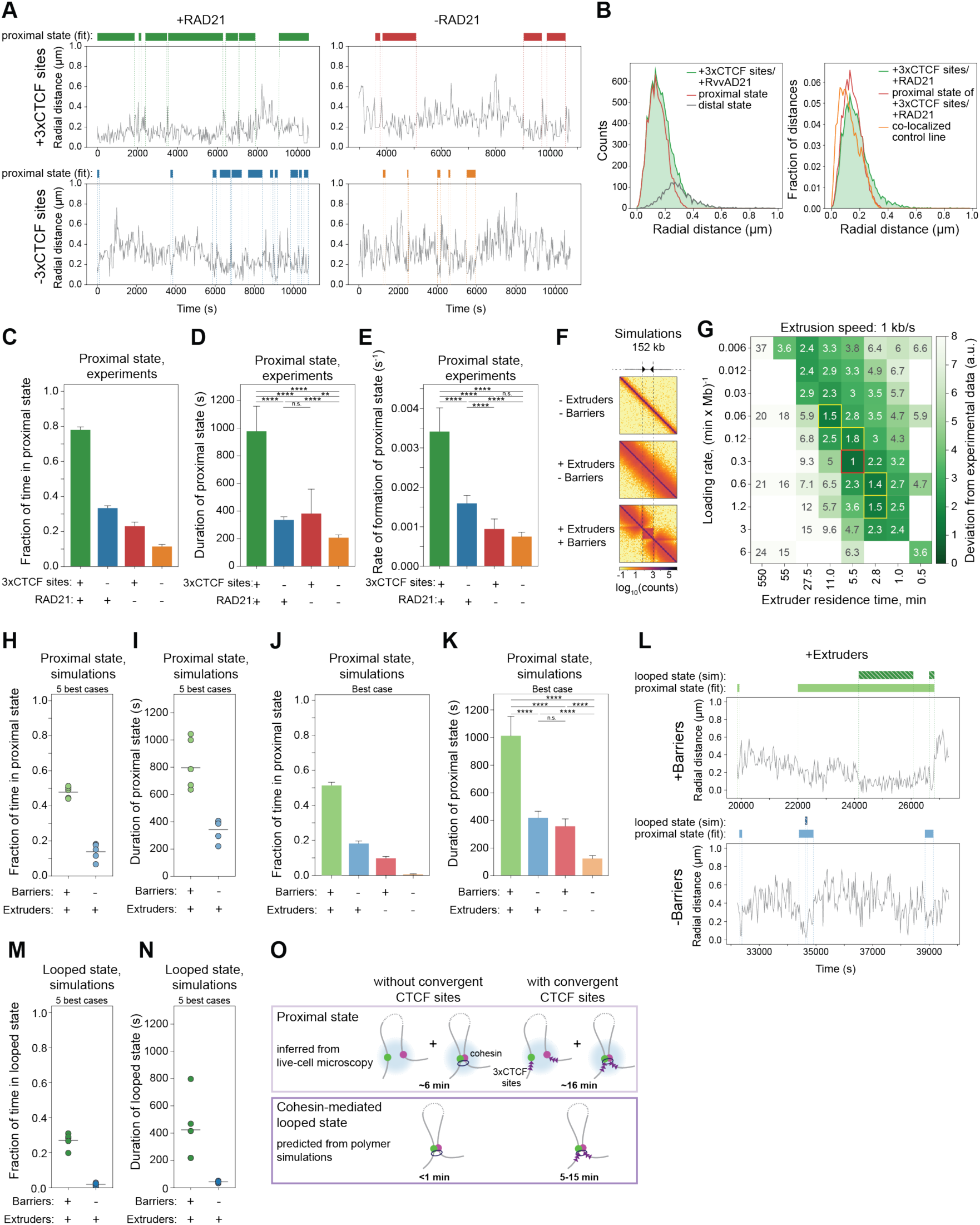
Cohesin and CTCF modulate the duration and rate of formation of contacts between sequences in *cis*. A. Representative trajectories of radial distance (gray) and occurrences of the proximal state called by HMM (colored bars). The HMM was fitted on data with convergent 3xCTCF sites and RAD21 (top left) and applied to the other three samples. B. Left: Radial distance distribution in cells with convergent 3xCTCF sites and RAD21 overlaid with those of proximal and distal states called by HMM on the same sample. Right: Same as in the left panel with the additional display of the distance distribution from a control cell line where TetO and LacO signals perfectly colocalize. C. Fraction of time spent in the proximal state called by HMM in the four experimental conditions (no. of replicates as indicated in Fig. 4F). Error bars represent bootstrapped (n=10000) standard deviations. D. Left: Average duration of proximal states (mean +/- 95% confidence interval). P-values (Student t- test, two-sided): *=p<0.05, **=p<0.01, ***=p<0.001, ****=p<0.0001. P-values can be found in **Suppl. Table S2**. Right: Average rates of contact formation (time elapsed between the end of a proximal state and the beginning of the next). E. Simulated contact maps of a polymer chain either without loop extrusion, or with loop extruders and of convergent one-sided impermeable extrusion barriers separated by approximately 150 kb. F. Levels of agreement between simulations and experimental data as a function of loop extrusion parameters (here shown with extrusion speed 1 kb/s). The score represents the deviation of the distance, duration and fraction of time spent in the proximal state with those experimentally observed in the presence of RAD21 with or without 3xCTCF sites (see Methods). Red square: parameter set maximizing the agreement with experimental values. Yellow squares: four additional second-best parameter sets. G. Fraction of time spent in the contact state called by HMM on simulations with best-matching parameters (red square in panel F for +Extruder case, see Methods). Error bars as in panel C. H. Duration (mean +/- 95% confidence interval) of contact state called by HMM on simulations with best-matching parameters. I. Changes in the average duration of contact states following introduction of RAD21 and convergent CTCF sites, or extruders and extrusion barriers in experiments and simulations, respectively. J. Representative trajectories of radial distances (gray), contact states called by HMM (full bar) and looped states in the underlying polymer conformations (striped bars) from +Extruders/+Barriers (top) and +Extruders/-Barriers simulations (bottom) with best-matching parameters (red square in panel F). K. Left: Fraction of time spent in the looped state based on simulations with best-matching parameters. Error bars as in panel C. Right: Durations of the looped state (mean +/- 95% confidence interval). P-values as in panel D. L. Scheme summarizing the durations of contact and looped states in the presence and absence of 3xCTCF sites.

We next fitted the HMM model to all experimental conditions while keeping the same proximal state as in cells with 3xCTCF sites and RAD21 (**Fig. 5A).** This showed that in the presence of RAD21, the LacO and TetO arrays spent approximately 78% of the time in contact (i.e. in the proximal state) when 3xCTCF sites were present. This was 2.4 times higher than the 33% of time they spent in contact when the 3xCTCF sites were removed (**Fig. 5C)**, in agreement with the corresponding 2.6-fold difference in contact probability inferred from Capture-C (**Fig. 4B**). The fraction of time spent in contact decreased markedly upon depletion of RAD21 to approximately 23% in the presence of 3xCTCF sites and 11% in the absence (**Fig. 5C**). Both the average duration of contacts and their rate of formation were maximal in the presence of RAD21 and convergent 3xCTCF sites, where they lasted around 16 minutes and reformed every 5 minutes on average (**Fig. 5D, E**). Contacts became substantially shorter (6 minutes) and rarer (one every 10 minutes on average) when 3xCTCF sites were removed, and even more so upon RAD21 depletion (lasting 2 minutes and occurring every 22 minutes on average). Thus both cohesin and CTCF impact both the duration and the probability of formation of chromosomal contact events between loci separated by 150 kb within an otherwise “empty” TAD.

To understand if these results could be rationalized in terms of the underlying loop extrusion process, we compared them to polymer simulations with convergent impermeable loop extrusion barriers separated by approx. 150 kb (**Fig. 5F**). Simulations were performed using loop extrusion parameters spanning a finer-grained 25-fold range around experimentally realistic values that reproduced the dynamic effect of RAD21 degradation (cf. **Fig. 2D,** black square: cohesin residence times of 5.5 minutes and linear densities of approx. 23 per Mb) and with both *in vitro* and *in vivo* estimates of extrusion speeds^19, 30^. To allow direct comparison with experimental distance-based HMM states, we applied random errors matching experimental uncertainty levels (see Methods) to radial distances generated by the models (**Suppl. Fig. S5C**) and called proximal and distal states using the same HMM strategy as with experimental data. Importantly, for a large number of parameter combinations, distances in the proximal state largely overlapped with the corresponding distribution observed experimentally in the presence of convergent CTCF sites and cohesin (**Suppl. Fig. S5D, Suppl. Fig. S6A**).

We then compared the distance, duration and fraction of time spent in the proximal state with those experimentally observed in the presence of RAD21 with or without 3xCTCF sites. We found that their similarity was maximal in a range of parameters corresponding to extruder densities of 8 to 32 per Mb (**Suppl. Fig. S6E**) and residence times of 2.8 to 11 minutes, with extrusion speeds of both 0.1 and 1 kb/s, all of which were in the range of previous estimations of experimental values^19, 30, 36^ (**Fig. 5G, Suppl. Fig. S5E**, see Methods). Considering the five best-matching scenarios (red and yellow marked values in **Fig. 5G**), the two locations spent 45% to 55% of the time in the proximal state with an average contact duration of around 10 to 17 minutes, which reduced to 18% and 8 minutes in the absence of extrusion barriers (**Fig. 5H,I**). In addition, similar to the effects observed experimentally upon depletion of RAD21, decreasing extruder densities (e.g. by decreasing loading rates) led to decreased durations and rate of formation of contacts (**Fig. 5J,K,** shown for the best case, red marked value in **Fig. 5G**, general trends in **Suppl. Fig. S6B-D,** see Methods). Thus solely relying on general estimates of realistic loop extrusion parameters and without any additional parameter fitting, a simple loop extrusion model with convergent barriers was able to reproduce the full range of dynamic properties observed in live-cell microscopy. This supports the notion that loop extrusion is largely responsible for our experimental observations.

The HMM-based proximal state likely provides an overestimation of the duration of underlying CTCF-CTCF loops mediated by stalled cohesins, since it also contains a fraction of CTCF-independent proximity events that cannot be distinguished from loops. To estimate the duration and time the two loci spent in a cohesin-mediated CTCF-CTCF looped conformation, we therefore quantified occurrences in the simulated polymer where the two monomers formed the base of an extruded loop, either in the presence or absence of extrusion barriers (**Fig. 5L**, see Methods). As expected, these events were rarer and shorter than contacts detected by HMM on polymer simulations (**Fig. 5H,I**), with two monomers spending approximately 20-31% of time at a loop base for 5-15 minutes on average in the presence of extrusion barriers (**Fig. 5M,N**). Finally, transient cohesin-dependent loops that are not stabilized by convergent CTCF sites should occur much more rarely (1-3% of the time) and last less than a minute on average (**Fig. 5M,N**). Comparison of actual configurations in the polymer simulations with HMM states thus suggests that the dynamics of chromosome contacts detected at a range of 150 nm is generated by faster and rarer cohesin-mediated CTCF loops (**Fig. 5O**).

## Discussion

Our study provides quantitative measurements of chromosome folding dynamics in living cells and reveals how it is controlled by cohesin and CTCF. Two experimental strategies allow us to minimize biological variation from specific regulatory and structural genomic contexts and enable direct comparison with polymer models. First, by studying large numbers of random genomic locations, we average over local differences in chromosome mobility and reveal the global dynamic effects of cohesin. Secondly, by visualizing and manipulating two locations within a TAD devoid of additional regulatory or structural features, we unravel how cohesin and CTCF impact chromosome looping at genomic distances that are representative of CTCF-mediated interactions and enhancer-promoter connections genome-wide. Our results reveal that although higher extrusion speeds could in principle result in an acceleration of chromosome motion (**Suppl. Fig. S5F,G**), physiological loop extrusion rates rather result in transient constraints that slow down chromosome dynamics. In line with previous reports^37^ we observe that constraints introduced by loop extrusion reduce the spatial distances between genomic sequences located in *cis*, thus increasing the chances that they meet in space. Our measurements now reveal that this entails an increase in both the rate of formation and the duration of contacts. Convergently oriented high-affinity CTCF motifs lead not only to higher contact frequencies as expected from Hi-C, but also substantially longer contact durations. This suggests that asymmetries in contact patterns established by CTCF motifs genome-wide might also lead to temporal asymmetries in physical interactions, notably between regulatory sequences. It might also provide a framework to understand how single CTCF sites can modulate gene expression, occasionally without noticeable changes in contact probabilities^3^. We additionally observe that constraints introduced by cohesin and convergent CTCF sites lead to reduced temporal variability in physical distances, arguing that loop extrusion increases the reproducibility of chromosome folding at selected genomic sites.

Finally, our study provides estimates for the frequency and duration of contacts defined by physical distances (∼150 nm) that might be comparable to those where signals arise in 3C methods^13, 17^. For sequences separated by 150kb inside the same TAD, such contacts assemble and disassemble on the timescale of minutes (although we cannot exclude that shorter-range contacts between DNA molecules and factors bound to them could be even faster). This provides many opportunities in a single cell cycle for regulatory sequences in a TAD to occur in close spatial proximity. Estimates based on comparison with polymer simulations further suggest that molecular configurations of the chromatin fiber where a loop is anchored by convergent CTCF sites located 150 kb apart might last around 5 to 15 minutes and occur around 27% of the time. This is in good agreement with recent estimates of the kinetics of a loop connecting the boundaries of the endogenous *Fbn2* TAD in mESC^21^, which however occurs more rarely (3.5-6% of time) possibly due to the longer distance involved (around 500 kb). Taken together, our data establish firm quantitative bases for understanding the dynamics of chromosome folding at the scale of TADs and provide temporal constraints for mechanistic models of chromosome structure and its impact on fundamental biological processes such as long-range transcriptional regulation.

## Acknowledgements

We would like to thank all members of the Giorgetti lab for help with labeling images for training the deepBlink spot-detection algorithm and feedback on the manuscript, Anders S. Hansen, Leonid Mirny and members of their labs for useful discussions of the results, Elzo de Wit for sharing the WAPL-AID-eGFP cells, and Hubertus Kohler for assistance with flow cytometry experiments. P.M. was supported by a Marie Sklodowska-Curie Innovative Training Network (813327 ‘ChromDesign’). J.T. was supported by a Marie Sklodowska-Curie Innovative Training Network (813282 ‘PEP-NET’). J.Z. was supported by a Marie Sklodowska-Curie grant (748091 - 3DQuant). Research in the Giorgetti lab is funded by the Novartis Foundation, the European Research Council (ERC) (759366, ‘BioMeTre’), Marie Sklodowska-Curie Innovative Training Networks (813327 ‘ChromDesign’ and 813282 ‘PEP-NET’) under the European Union’s Horizon 2020 research and innovation program, and the Swiss National Science Foundation (310030_192642).

## Author contributions

PM and PK conceived the study with LG and with the help of discussions with EPN. PM performed the experiments with help from JT, JC, SG, JZ, MK. PK developed the code, performed and analyzed polymer simulations with input from EM and GT. YZ performed image and trajectory analysis with help from PK, EM and GT. LGe and JE contributed setting up and provided assistance with microscopy and image analysis. SS performed piggyBac insertion site mapping and provided assistance with high-throughput sequencing experiments. PK and YZ analyzed sequencing data. LG wrote the paper with PM, PK, YZ, GT and input from all the authors.

---

**Supplementary Figure S1:**
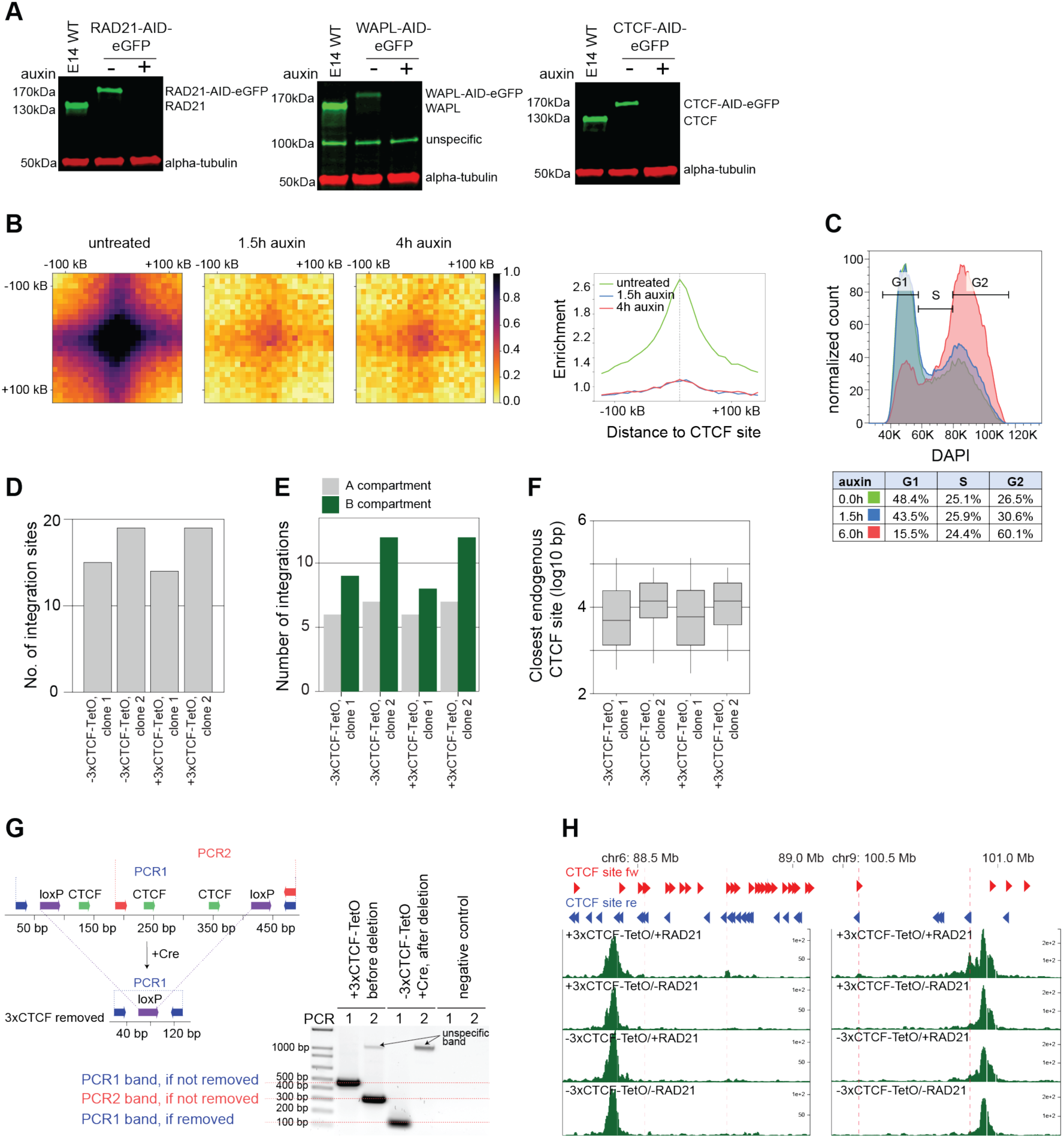
Chromosome structure is altered upon degradation of factors involved in loop extrusion. A. Western Blots showing the degradation upon treatment with 500 µM auxin of either RAD21 after 1.5 h (left), WAPL after 24 h (middle) or CTCF after 6 h (right). Loading control: *α*-tubulin. B. Left: Pile-up plots of enrichment in contact probability at CTCF sites based on Hi-C data in RAD21- AID-eGFP cells that were either untreated (left), treated for 1.5 h (middle) or 4 h (right) with 500 µM auxin to induce degradation of RAD21. Right: Differences in contact enrichment at CTCF peaks. Peaks were called on Hi-C data from untreated cells. C. Flow cytometry analysis of fixed cells stained with 4′,6-diamidino-2-phenylindole (DAPI) showing the cell-cycle stage distribution of RAD21-AID-eGFP mESC cultured in the presence of serum, leukemia inhibitory factor (LIF) and 2i inhibitors. Cells were either untreated (green) or treated with 500 µM auxin for 1.5 h (blue) and 6 h (red). D. Integration site mapping of two clones from RAD21-AID-eGFP cell lines with (+3xCTCF-TetO) and without (-3xCTCF-TetO) 3xCTCF sites. E. Distribution of integration sites across A and B compartments, called on distance-normalized Hi-C map using 1st eigenvector and matched with epigenetic marks, for the same clonal cell lines also shown in panel E. F. Distance distribution of integration sites shown in panel E from the closest endogenous CTCF sites (endogenous CTCF sites from^9^). Boxplot: boxes denote lower and upper quartiles (Q1 and Q3, respectively); whiskers denote 1.5× the interquartile region (IQR) below Q1 and above Q3. G. Example of genotyping PCR upon the removal of 3xCTCF sites in one RAD21-AID-eGFP + 3xCTCF-TetO clonal cell line. PCR1 amplifies the entire 3xCTCF cassette and the product size changes from 470 bp to 147 bp if the cassettes are successfully removed. PCR2 amplifies half of the 3xCTCF cassette and no product is expected if 3xCTCF cassettes were removed from all insertion sites; otherwise a PCR band of 303 bp is expected. H. Representative 4C-seq profiles from insertions on chromosomes 6 and 9 using TetO as a viewpoint showing that 3xCTCF-TetO cassettes lead to the formation of ectopic contacts (dashed red lines) with nearby endogenous CTCF sites in the presence of RAD21. These contacts are lost upon deletion of the 3xCTCF cassette (-3xCTCF-TetO) and upon degradation of RAD21 (-RAD21).

**Supplementary Figure S2:**
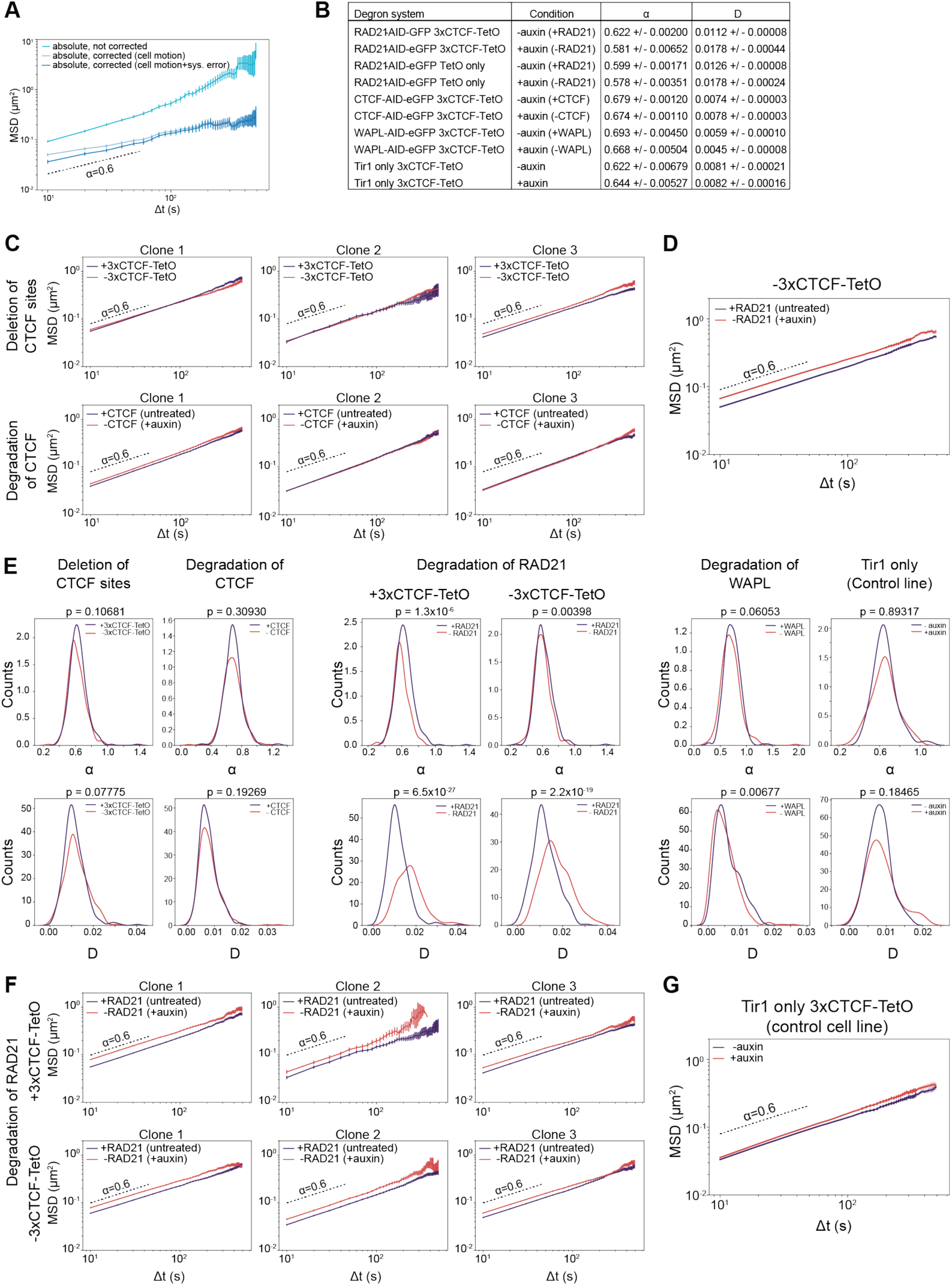
Chromosome dynamics is modulated by degradation of factors involved in loop extrusion. A. Mean Square Displacement (MSD) of trajectories from TetO insertions within the same cell (MSD, mean +/- standard error) before (cyan) and after applying cell motion (light blue) and localisation error correction (dark blue). B. Scaling exponents (*α*) and generalized diffusion coefficients (D) across all conditions and cell lines were fitted by pooling all the three biological replicates. Shown are the numbers for the best fit +/- error of the fit. C. MSD (mean +/- standard error) plots for a single clonal cell line (biological replicate) when looking at removal of 3xCTCF sites (top row) next to the array or degrading all CTCF (bottom row). D. MSD (mean +/- standard error) in the cell lines (n=3 replicates per clonal cell line, three cell lines) where the 3xCTCF cassette was excised. Shown are the MSDs for cells either depleted of RAD21 for 90 min (red, 266 cells, 9,020 trajectories analyzed) or not (blue, 271 cells, 11,082 trajectories analyzed). Global depletion of RAD21 increases mobility. P-values in panel E. E. Distributions of *α* and D fitted based on single trajectory MSD and significance test for differences in generalized diffusion coefficients (D) and scaling exponents (*α*). The p-value is calculated using Student t-test (two-sided) (see Methods). F. Same as in C for a single clonal cell line (biological replicate) with integrations with 3xCTCF-TetO (top row) or without 3xCTCF-TetO (bottom row) when degrading RAD21. Global depletion of RAD21 increases mobility. G. Same as in D in the cell lines that contain integrations of 3xCTCF-TetO and the Tir1 protein, but do not contain any AID-tag for targeted degradation. MSDs for cells either treated with auxin for 90 min (red, 97 cells, 2,155 trajectories analyzed) or not (blue, 111 cells, 3,711 trajectories analyzed). No significant changes were detected. P-values in panel E.

**Supplementary Figure S3:**
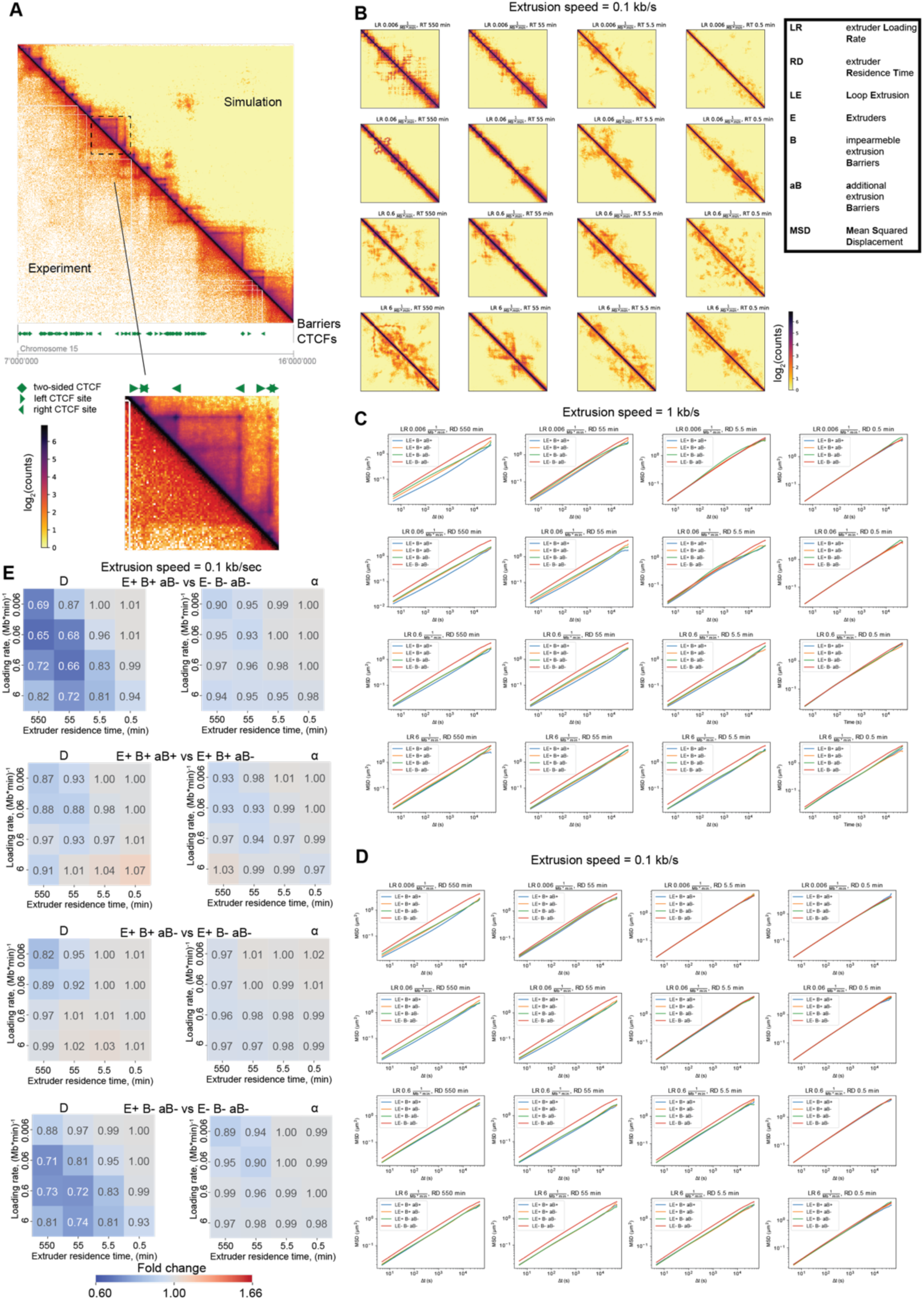
Chromosome dynamics are modulated by loop extrusion. A. Visual comparison of experimental Hi-C contact map with contact maps of simulations at extrusion speed 1 kb/s, extruder loading rate 0.06 (Mb x min)^^-1^ and residence time 5.5 min. B. Contact maps for the polymer simulations at extrusion speed 0.1 kb/s and barriers from the range 7-16 Mb of chromosome 15. Acronyms used in this figure are indicated in the black box on the right. C. MSDs for all 4 conditions for each set of loop extrusion parameters and extrusion speed of 1 kb/s. D. Same as C but for the extrusion speed of 0.1 kb/s. E. Pairwise comparison for conditions indicated in the title of each pair of heatmaps. Pair of heatmaps contains ratios of generalized diffusion coefficients (D) and scaling exponent (*α*), and represents fold change between the conditions.

**Supplementary Figure S4:**
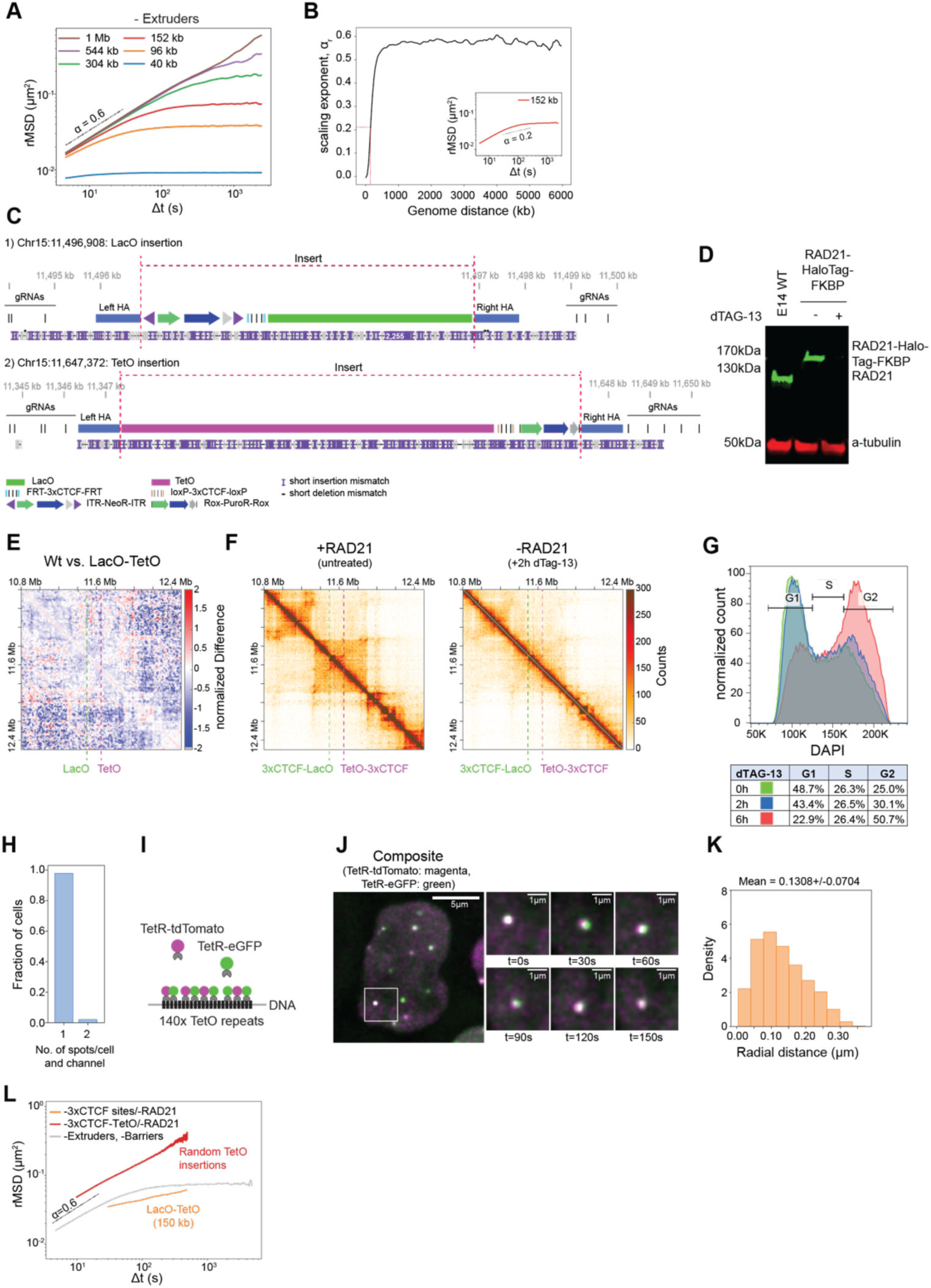
DNA loci in *cis* separated by 150 kb show a more correlated motion that is modulated by CTCF and cohesin. A. rMSD of multiple pairs of monomers separated by various distances 40 kb - 1 Mb. Simulations were performed for the polymer without extruders and barriers. Values were averaged with a sliding window without considering the first and last 200 monomers (1.6 Mb). Dashed scaling exponent *α*=0.6 serves as an eyeguide. B. Distance dependency of the scaling exponent (*α*) on the genomic distance between loci. C. IGV snapshot showing an example of a Nanopore sequencing read mapped to a modified mouse genome including the respective insertions. Reads that spanned from a guide RNA (gRNA) binding site upstream of the left homology arm (left HA) to a gRNA binding site downstream the right homology arm (right HA) confirmed single insertion of the transgene. D. Western Blots showing the targeted degradation of RAD21 after 2 h of treatment with 500 nM dTAG-13. Loading control: anti-tubulin. E. Differential map at 6.4 kb resolution for the structural differences between a E14 wild-type (Wt) and the E14 cell line containing LacO and TetO insertions (see Methods). Dashed lines indicate the insertion sites. No structural changes are detected upon integration of the operator arrays. F. Capture C maps at 6.4kb resolution in the region on chr15 (10.8 Mb-12.5 Mb) in the untreated cells (left) and in cells treated with 500 nM dTag-13 (left) showing that RAD21 degradation leads to loss of chromosome structure. G. Flow cytometry analysis of fixed cells stained with DAPI to show cell cycle stage distribution of E14 RAD21-HaloTag-FKBP cells. H. Bar plot showing the number of detected spots per cell per channel for 1,400 manually annotated images subsampled from the images series. In 3% of the images 2 spots per cell are detected indicating the presence of sister-chromatids. I. Schematic representation of the control cell line that contains multiple TetO array integrations as well as stable integrations of the TetR-eGFP and TetR-tdTomato expressing cassettes. This allowed the labeling of the TetO with two separate fluorophores. J. Representative images of the control cell line. The time series shows a zoomed version of the region indicated by the white square. K. Radial distance distribution of the control cell line showing that the resolution on the 3D distance is approximately 130 nm. L. rMSD for cell lines containing multiple random integrations of -3xCTCF-TetO as shown in Fig. 1E (cyan) or the targeted integrations of LacO and TetO on chr15 (orange) in the absence of RAD21 compared to the predicted rMSD of two loci at a distance of 150 kb in the absence of extruders (gray) as predicted from polymer simulations.

**Supplementary Figure S5:**
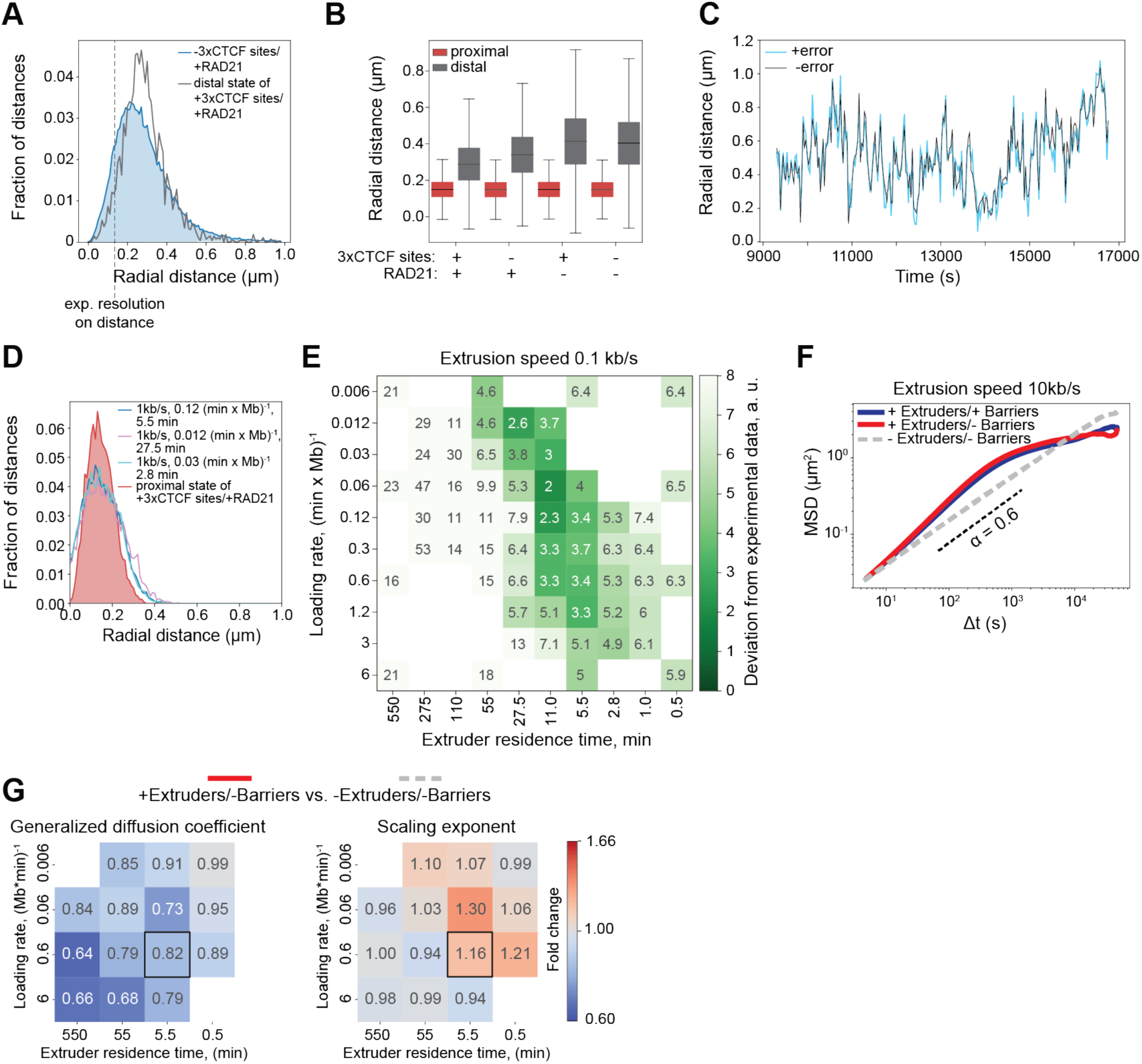
A. Radial distance distribution for the condition -3xCTCF sites/+RAD21 (blue) overlaid with the noncontact state called by HMM on the +3xCTCF sites/+RAD21 (gray) showing that the noncontact state identified by HMM largely overlaps with the distance distribution of the two loci in the absence of the CTCF sites. B. Boxplot for the radial distances for the contact and noncontact state called by HMM on all four conditions. The horizontal line indicates the median. Box plots are as in **Suppl. Fig. S1F**. C. Representative radial distance trajectory of a simulated system with and without an additional error on the distance that is in the range of the experimental error. D. Radial distance distribution for the contact state of the +3xCTCF sites/+RAD21 condition overlaid with the distributions of the contact states from the three best matching parameters sets when comparing only the average radial distances. E. Heatmap showing the agreement of all simulated systems (for extrusion speed 0.1 kb/s) with the experimental data. The score is as described in Fig. 5F (see Methods). F. MSDs for 3 conditions for extruder residence time of 5.5 min, loading rate of 0.6 Mb x min^-1^ and extrusion speed of 10 kb/s. Pairwise comparison for conditions indicated in the title of each pair of heatmaps. G. Heatmap showing the fold change of generalized diffusion coefficients (D) and scaling exponent (*α*), and represents fold change between the conditions.

**Supplementary Figure S6:**
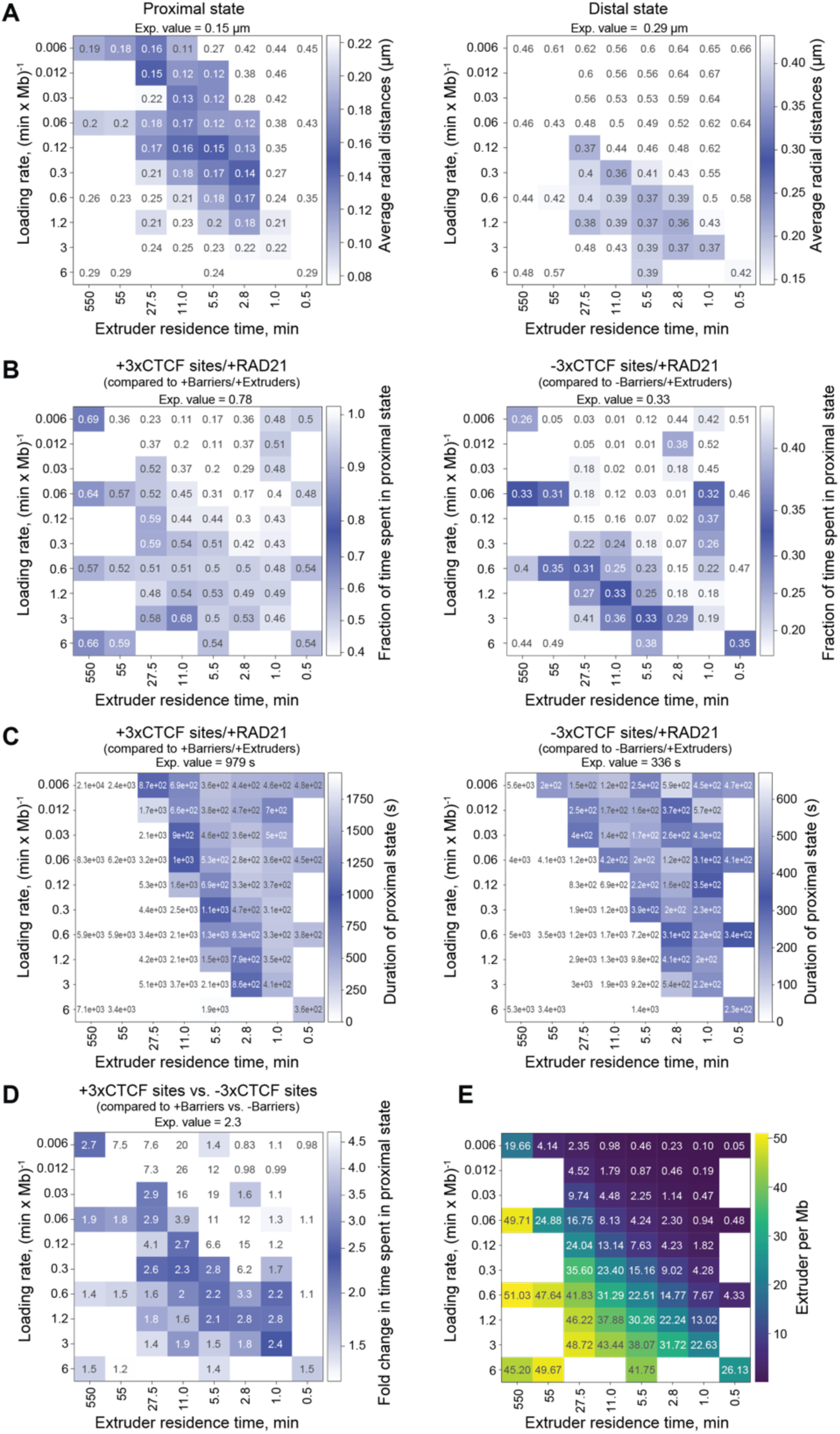
A. Heatmap showing the agreement of average radial distances of either the contact state (left) or the noncontact state (right) called by HMM on all simulated systems of extrusion speed 1 kb/s with the contact state or noncontact state called by HMM on the experimental data. Darker shades of blue indicate better agreement with experimental values. B. Same as in panel A, but for the fraction of time spent in the contact state (left) and noncontact state (right). C. Same as in panel A but for the duration of the contact state for either the condition +Barriers/+3xCTCF sites (left) or -Barriers/-3xCTCF sites (right) in the presence of the Extruders/RAD21. D. Same as in panel A but for the fold change of time spent in a contact state called by HMM on the experimental data for the comparison of the condition +Barriers/+3xCTCF sites vs. -Barriers/- 3xCTCF sites in the presence of Extruders/RAD21. E. Extruder densities per Mb for all simulated systems with extrusion speed 1 kb/s. Color-coded for the density per Mb.

## SUPPLEMENTARY VIDEO LEGENDS

**Supplementary Video S1: Live-cell imaging od TetO arrays upon depletion of RAD21**

Time course of RAD21 degradation upon induction with 500 µM auxin in RAD21-AID-eGFP cells. TetO integrations are tagged with TetR-tdTomato (magenta) and RAD21 is tagged with eGFP (green). Green fluorescence is lost within 90 min after induction of degradation (exposure time (eGFP) = 50 ms, exposure time (tdTomato) = 50 ms, deconvolved, max. intensity projection, duration of movie 30 min, dt=10 s).

**Supplementary Video S2: Dynamics of LacO-TetO radial distances**

Representative movie of dual-color imaging of LacO (green) and TetO (magenta) arrays flanked by 3xCTCF sites integrated on chromosome 15 at a distance of 150 kb (exposure time (eGFP) = 50 ms, exposure time (tdTomato) = 50 ms, deconvolved, max. intensity projection, duration of movie 1 h, dt=30 s).

**Supplementary Video S3: Cohesin and CTCF decrease average LacO-TetO radial distances**

Representative movies of dual-color imaging of LacO (green) and TetO (magenta) arrays on chromosome 15 at a distance of 150 kb. Left panel: Cell line with 3xCTCF sites flanking LacO and TetO (in the presence of RAD21); middle panel: Cell line where 3xCTCF sites have been removed (in the presence of RAD21); Right panel: Cell line with 3xCTCF sites flanking the array, but where RAD21 has been degraded with 500 nM dTAG-13 (exposure time (eGFP) = 50 ms, exposure time (tdTomato) = 50 ms, deconvolved, max. intensity projection, duration of movie 1 h, dt=30 s).

## METHODS

### Culture of embryonic stem cells

All cell lines are based on E14 mouse embryonic stem cells (mESCs). WAPL-AID-eGFP mESCs^38^ were kindly provided by Elzo de Wit (Netherlands Cancer Institute). All cell lines for the dual array imaging approach are based on the double-CTCF knockout cell line described in Zuin et al.^3^. Cells were cultured on gelatin-coated culture plates in Glasgow Minimum Essential Medium (Sigma-Aldrich, G5154) supplemented with 15% foetal calf serum (Eurobio Abcys), 1% L-Glutamine (Thermo Fisher Scientific, 25030024), 1% Sodium Pyruvate MEM (Thermo Fisher Scientific, 11360039), 1% MEM Non-Essential Amino Acids (Thermo Fisher Scientific, 11140035), 100 µM β-mercaptoethanol (Thermo Fisher Scientific, 31350010), 20 U/ml leukemia inhibitory factor (Miltenyi Biotec, premium grade) in 8% CO2 at 37°C. Cells were tested for mycoplasma contamination regularly and no contamination was detected. After genome engineering and for Hi-C, Capture-C, 4C-Seq, Western Blot and imaging experiments, cells were cultured in standard E14 medium supplemented with 2i (1 µM MEK inhibitor PDO35901 (Axon, 1408) and 3 µM GSK3 inhibitor CHIR 99021 (Axon, 1386)). For live-cell imaging experiments, cells were cultured in Fluorobrite Dulbecco’s Modified Eagle Medium (DMEM) (Gibco, A1896701) supplemented with 15% foetal calf serum (Eurobio Abcys), 1% L-Glutamine (Thermo Fisher Scientific, 25030024), 1% Sodium Pyruvate MEM (Thermo Fisher Scientific, 11360039), 1% MEM Non-Essential Amino Acids (Thermo Fisher Scientific, 11140035), 100 µM β-mercaptoethanol (Thermo Fisher Scientific, 31350010), 20 U/ml leukemia inhibitory factor (Miltenyi Biotec, premium grade) and with 2i inhibitors (1 µM MEK inhibitor PDO35901 (Axon, 1408) and 3 µM GSK3 inhibitor CHIR 99021 (Axon, 1386)).

### Generation of targeting vectors for random integration of TetO array and TetR-tdTomato

To generate a 8 kb TetO array within piggyBac ITRs (inverted terminal repeats), the TetO array was obtained from the pSO2.Pac.TetO, a gift from Edith’s Heard lab^39^, by growing bacteria at 37°C to reduce the size of original 30 kb operator array by recombining. The array was then excised from the vector by restriction digest with BamHI (NEB, R0136S) and cloned into the PB-empty^40^ (final vector PB-empty-DSE- TetO-8kb). A cassette carrying three strong CTCF sites was excised with XhoI (NEB, R0146S) from PB- empty_DSE_TetO_2.7kb_3xCTCF^40^) and ligated into the PB-empty-DSE-TetO-8kb vector using T4 DNA Ligase (NEB, M0202L). Clones were screened by Sanger sequencing (Microsynth) for CTCF sites inserted facing towards the TetO array. The final vector (PB-3xCTCF-TetO) was validated by restriction digest with EcoRI (NEB, R3101L) and NotI (NEB, R3189L) for correct size of the operator array and Sanger sequencing (Microsynth) for correct insertion of the CTCF cassette.

To express the Tet repressor (TetR) and Lac repressor (LacI) fused to a fluorescent protein flanked by ITRs for piggyBac transposition, the PB-empty vector was first linearized by digestion with XhoI (NEB, R0146L) and the TetR-eGFP was amplified with Phusion High-Fidelity DNA Polymerase (Thermo Fisher Scientific,

F530L) from pBroad3-TetR-ICP22-EGFP kindly provided by Tim Pollex^41^ with Gibson overhangs. PB-Ubc- TetR-eGFP was assembled using Gibson cloning (NEB, E2611L). The tdTomato was amplified with Gibson overhangs and assembled with the digested PB-TetR-eGFP (BamHI, NEB, R3136L and EcoRI, NEB, R3101L) to yield the PB-Ubc-TetR-tdTomato vector. To increase expression levels of the fusion proteins, the Ubc promoter was exchanged for the stronger CAGGS promoter that was amplified with Gibson overhangs from a pCAGGS plasmid. The final PB-TetR-tdTomato was made using Gibson assembly of the amplified CAGGS promoter (from Addgene plasmid #20733) with the PB-Ubc-TetR-tdTomato digested with BglII (NEB, R0144L) and AgeI (NEB, R3552S). To generate PB-CAGGS-TetR-eGFP, PB-Ubc-TetR-eGFP and PB-TetR-tdTomato were digested with XhoI (NEB, R0146L) and ligated using T4 DNA Ligase (NEB, M0202L). PB-LacI-eGFP was generated by amplification of the LacI with overhangs for subsequent Gibson assembly with the digested PB-TetR-eGFP (AgeI, NEBR3552S). Primers used for cloning can be found in **Suppl. Table S3.**

### Generation of targeting vectors for TetO and LacO

Vector for targeting the TetO array to the genomic locus on chr15:11,647,372: The vector pMK-chr15-Rox- PuroR-Rox containing the homology arms for chromosome 15 as well as the Puromycin resistance gene flanked by Rox sites was custom synthesized by GeneArt Synthesis (Thermo Fisher Scientific) and linearized with SbfI (NEB, R0642S) and SpeI (NEB, R3133S). A short linker sequence including a XhoI restriction site was introduced into the vector by PCR amplification from pMK-chr15-Rox-PuroR-Rox with Gibson overhangs. The XhoI restriction site was then used to ligate the 3xCTCF-TetO (cut from the PB- 3xCTCF-TetO vector) into the vector leading to pMK-3xCTCF-TetO-Rox-PuroR-Rox.

Vector for targeting the LacO array to the genomic locus on chr15:11,496,908: The vector pUC19-empty was linearized with EcoRI (NEB, R3101L) and BamHI (NEB, R3136L). The left homology arm was amplified from E14 wild-type genomic DNA with overhangs for Gibson assembly. The 5’-ITR was amplified from PB-empty with Gibson overhangs and both PCR products were assembled into the pUC19 vector (Addgene, #50005) to yield pUC19-lHA-5’ITR. pUC19 was digested with KpnI (NEB, R3142L) and BamHI (NEB, R2126L) to be assembled into pUC19-3’ITR-3xCTCF-rHA with the following PCR products with respective Gibson overhangs: The 3’ITR was amplified from PB-empty, the CTCF cassette was amplified from pMK- chr15-Rox-PuroR-Rox and the right homology arm was amplified from E14 wild-type genomic DNA. To make the targeting vector, pUC19-lHA-5’ITR was linearized with EcoRI and BamHI, pUC19-3’ITR-3xCTCF- rHA was linearized with KpnI and HindIII (NEB, R3104L) and the Neomycin resistance gene was amplified from pEN113 (Addgene, #86233). All three parts were assembled using Gibson assembly to pUC19-ITR- NeoR-ITR-3xCTCF. The LacO array was excised from the pLAU43_LacO_plus vector^42^ with XhoI (NEB, R0146L) and ligated into the linearized targeting vector (cut with XhoI) using T4 DNA Ligase resulting in the final targeting vector pUC19-ITR-NeoR-ITR-3xCTCF-LacO. Primers used for cloning can be found in **Suppl. Table S3.**

### Generation of mESC lines carrying random integrations of TetO arrays

To generate clonal cell lines carrying random integrations of the TetO array in the degron cell lines (E14 Rad-AID-eGFP, E14 CTCF-AID-eGFP and E14 WAPL-AID-EGFP) 0.5×10^6^ cells were transfected with 2 µg PB-3xCTCF-TetO vector, 200 ng PB-TetR-tdTomato and 200 ng pBroad3_hyPBase_IRES_tagRFPt^40^ with Lipofectamine3000 (Thermo Fisher Scientific, L3000008) according to the manufacturer’s recommendations. Cells were cultured in standard E14 medium for 5 days and subsequently sorted by fluorescence-activated cell sorting (FACS sort) for fluorescent emission at 581 nm (tdTomato) on 96 well-plate to isolate clonal lines. Sorted cells were kept for 2 days in standard E14 medium supplemented by 100 μg/µl primorcin (InvivoGen, ant-pm-1) and 10 µM ROCK inhibitor (STEMCELL Technologies, Y- 27632). 10 days after sorting the plates were duplicated by detaching with accutase (Sigma Aldrich, A6964) and re-seeding in full E14 culture medium. 1/3 of the cells were replated onto Corning High-Content Imaging Glass Bottom Microplates (96-well, Corning, 4580). 2 days after reseeding, clonal lines were screened by microscopy for >10 insertions of TetO/cell and a good signal-to-noise ratio (SNR). Selected clones were expanded and genotyped by PCR for absence of random integration of the PiggyBase itself. Primers used for genotyping are listed in **Suppl. Table S3.**

### Removal of CTCF sites by Cre recombination

To selectively remove the three CTCF binding sites flanking the operator arrays, 0.5×10^6^ cells of each cell line were transfected 1 µg of pIC-Cre (a gift from the Schübeler lab) using Lipofectamine3000 according to the manufacturer’s instructions (Thermo Fisher Scientific, L3000008). 7 days after transfection, the cells were sorted and genotyped as described previously. Primers used in genotyping are listed in **Suppl. Table S3.**

### Generation of control cell lines expressing OsTir1

E14 wild-type cells were transfected with the targeting vector pEN396-pCAGGS-Tir1-V5-2A-PuroR TIGRE donor and the gRNA vector pX330-EN1201 (Addgene #92144 and #92142) using nucleofection with the Amaxa 4D-Nucleofector X-Unit and the P3 Primary Cell 4D-Nucleofector X Kit (Lonza, V4XP-3024 KT). 2×10^6^ cells were harvested using accutase (Sigma Aldrich, A6964) and resuspended in 100 µl transfection solution (82 µl primary solution, 18 µl supplement, 15 μg Tir1 targeting vector and 5 µg of pX330-EN1201) and transferred to a single Nucleocuvette (Lonza). Nucleofection was performed using the protocol CG110. Transfected cells were directly seeded in pre-warmed E14 standard medium. 48 hours after transfection, 1 µg/ml of puromycin (InvivoGen, ant-pr-1) was added to the medium for 3 days to select cells for insertion of the Tir1 integration. Cells were sorted and genotyped as described previously. Primers used in genotyping are listed in **Suppl. Table S3.**

### Generation of dual array (TetO-LacO) mESC line

Integration of the TetO array into the genomic locus on chr15:11,647,372: The vector containing the gRNA sequence was available from a previous study (PX459-chr15_gRNA/Cas9^3^). The gRNA sequence can be found in **Suppl. Table S3.** E14 mESC already containing a double-knockout for CTCF sites (clone D6 in Ref. ^3^) were transfected with the targeting vector pMK-3xCTCF-TetO-Rox-PuroR-Rox and the gRNA vector PX459-chr15_gRNA/Cas9 using nucleofection with the Amaxa 4D-Nucleofector X-Unit and the P3 Primary Cell 4D-Nucleofector X Kit (Lonza, V4XP-3024 KT). 2×10^6^ cells were nucleofected with 1 μg TetO targeting vector and 1 µg of PX459-ch15_gRNA/Cas9) as described above and treated with 1 µg/ml of puromycin (InvivoGen, ant-pr-1) 48h after transfection for 3 days to select cells for insertion of the TetO cassette. Cells were then cultured in standard E14 medium for additional 7 days and subsequently sorted by fluorescence-activated cell sorting (FACS sort) on 96 well-plate as described above to isolate clonal lines. 10 days after sorting the plates were duplicated by detaching with accutase (Sigma Aldrich, A6964) and re-seeding in full E14 culture medium. Genomic DNA was extracted on plate by lysing cells with lysis buffer (100 mM Tris-HCl pH 8.0, 5 mM EDTA, 0.2% SDS, 50 mM NaCl and 1 mg/ml proteinase K (Macherey-Nagel, 740506) and 0.05 mg/ml RNase A (Thermo Fisher Scientific, EN0531) and subsequent isopropanol precipitation. Individual cell lines were analyzed by genotyping PCR to determine heterozygous insertion of the TetO cassette. Cell lines showing the corrected genotype were selected and expanded. Primers used for genotyping are listed in **Suppl. Table S3.** Targeted nanopore sequencing with Cas9-guided adapter ligation^34^ (as described below) was performed on expanded clones to confirm single-copy insertion of the TetO cassette. Clone 2G5 was used for further engineering.

Integration of the LacO array into the genomic locus on chr15:11,496,908: The gRNA sequence for the CRISPR/Cas9 knock-in of the LacO cassette was designed using the online tool https://eu.idtdna.com/site/order/designtool/index/CRISPR_SEQUENCE and purchased from Microsynth AG. The gRNA sequence can be found in **Suppl. Table S3.** The gRNA sequence was cloned into the PX330 plasmid (Addgene, #58778) using the BsaI restriction site. The clonal line carrying the TetO cassette (clone 2G5) was transfected with the targeting vector pUC19-ITR-NeoR-ITR-3xCTCF-LacO and the gRNA vector pX330-chr15_LacO_gRNA/Cas9 using nucleofection with the Amaxa 4D-Nucleofector X-Unit and the P3 Primary Cell 4D-Nucleofector X Kit (Lonza, V4XP-3024 KT) as described for the Tir1 integration. 48 hours after transfection, 250 µg/ml of G418 (InvivoGen, ant-gn-1) was added to the medium for 3 days to select cells for insertion of the LacO cassette. Cells were sorted and genotyped as described for the TetO integration. Primers used for genotyping are listed in **Suppl. Table S3.** Cell lines showing the corrected genotype were selected and expanded. Expanded clones were transiently transfected with 200 ng PB- TetR-tdTomato and 200 ng PB-LacI-eGFP using Lipofectamine3000 according to the manufacturer’s instructions (Thermo Fisher Scientific, L3000008) and 2 days after transfection validated for heterozygous insertion of the LacO cassette on the same allele as the TetO by microscopy. Targeted nanopore sequencing with Cas9-guided adapter ligation^34^ (as described below) was performed on correct clones to confirm single-copy insertion of the LacO cassette. Clone 1F11 was used for further engineering.

### Integration of TetR-tdTomato and LacI-eGFP and removal of Puromycin resistance

To visualize the operator arrays in live-cell imaging and remove the puromycin resistance gene used for selection during integration, 0.5×10^6^ E14 TetO-LacO cells (clone 1F11) were transfected with 200ng PB- TetR-tdTomato, 200ng PB-LacI-eGFP and 200ng pBroad3_hyPBase_IRES_tagRFPt^40^ with Lipofectamine3000 (Thermo Fisher Scientific, L3000008) according to the manufacturer’s instructions. 7 days after transfection the cells were sorted (as described previously) for fluorescent emission at 507 nm (eGFP) and 581 nm (tdTomato). Sorted cells were cultured and genotyped as described for the random TetO integration. Primers used for genotyping are listed in **Suppl. Table S3.** Cell lines showing the corrected genotyping pattern were selected and expanded and a good and comparable SNR was selected for by microscopy. Clone 2C10 was used for further engineering.

### Tagging of endogenous *Rad21* locus and removal of resistance genes NeoR and HygroR

The gRNA vector and the targeting vector from Liu et al.^25^ were purchased from Addgene (gRNA pX330- EN1082: #156450, targeting vector pEN313: #156431). The gRNA sequence can be found in **Suppl. Table S3.** The targeting vector was modified using a restriction digest with NdeI and EcoRI (NEB, R0111S and NEB, R3101L) and subsequent Gibson assembly (NEB, E2611L) to insert an FKBP-tag at the end of the coding sequence as well as a Rox-HygroR-Rox cassette for selection (final vector: RAD21-Halo-FKBP- Rox-HygroR-Rox). The clonal lines carrying the TetO and LacO cassettes as well as TeR-tdTomato and LacI-eGFP (clone 2C10) were transfected with the targeting vector (RAD21-Halo-FKBP-Rox-HygroR-Rox) and the gRNA vector PX330-EN1082 using nucleofection with the Amaxa 4D-Nucleofector X-Unit and the P3 Primary Cell 4D-Nucleofector X Kit (Lonza, V4XP-3024 KT) as described above. 48 hours after transfection, 160 µg/ml of Hygromycin B (Thermo Fisher Scientific, 10687010) was added to the medium for 3 days to select cells for insertion of the HaloTag-FKBP-tag cassette. 7 days after selection, 0.5×10^6^ cells were transfected with 2 µg of pCAGGS-Dre-IRES-bsd (purchased from Gene Bridges) using Lipofectamine3000 according to the manufacturer’s instructions (Thermo Fisher Scientific, L3000008) to remove the Neomycin and Hygromycin resistances used as selection markers for previous integrations. Prior to single-cell sorting, the cells were incubated with 100 nM JF646 HaloTag Ligand^43^ in full culturing medium for 30 min at 37°C, 8%CO2 and washed three times with PBS. The cells were sorted for fluorescent emission at 664 nm. Sorted cells were cultured and genotyped as described for the Tir1 integration. Primers used for genotyping are listed in **Suppl. Table S3.** Cell lines showing the corrected genotype were selected, expanded and validated for homozygous insertion of the HaloTag-FKBP-tag and correct functioning of the degron system by Western Blot (as described below). Clone 1B1 was used for further engineering.

### Removal of CTCF sites in dual array TetO-LacO cell lines

To selectively remove the three CTCF binding sites flanking the operator arrays, 0.5×10^6^ cells of the TetO- LacO dual array cell line + RAD21-HaloTag-FKBP (clone 1B1) were transfected 1 µg of pIC-Cre using Lipofectamine3000 according to the manufacturer’s instructions (Thermo Fisher Scientific, L3000008). 5 days after transfection, 0.5×10^6^ cells of the pool were further transfected with 1 µg pCAG-FlpO-P2A- HygroR^3^. Cells were then sorted and genotyped as described previously. Primers used for genotyping are listed in **Suppl. Table S3.** Since the recombination with Flippase did not work sufficiently to attain correct clones, the CTCF sites flanking the LacO array were then removed using CRISPR-Cas9 deletion. The gRNA sequence for the CRISPR/Cas9 knock-out of the CTCF sites flanking the LacO array were designed using the online tool https://eu.idtdna.com/site/order/designtool/index/CRISPR_SEQUENCE and purchased from Microsynth. The gRNA sequence can be found in **Suppl. Table S3.** The gRNA sequence was cloned into the PX330 plasmid (Addgene, #58778) using the BsaI restriction site as described previously. The pool of cells transfected with pIC-Cre was further transfected with 0.5 µg of each gRNA/Cas9 vector and subsequently sorted and genotyped. Correct clones were expanded, validated by genotyping PCR that was analyzed on a 1% agarose gel imaged with a Typhoon FLA 9500 scanner (GE Healthcare). Subsequent Sanger sequencing of the PCR product (Microsynth) confirmed the removal and clones were used in live-cell imaging and capture-C experiments.

### Generation of control cell lines to measure localization error and experimental uncertainty

To generate a control cell lines for the dual array imaging, one clone of RAD21-AID-eGFP + 3xCTCF- TetO + TetR-tdTomato (clone 2B10) was transfected with 200 ng pBroad3_hyPBase_IRES_tagRFPt and 200 ng PB-TetR-eGFP using Lipofectamine3000 (Thermo Fisher Scientific, L3000008) according to the manufacturer’s instructions. Cells were cultured in standard E14 medium for 7 days and sorted (as described previously) for fluorescent emission at 507 nm (eGFP) and 581 nm (tdTomato). Clonal lines were screened for a good SNR by microscopy on Corning High-Content Imaging Glass Bottom Microplates (96- well, Corning, 4580) and were used for estimation of localisation error and the experimental uncertainty on the distance by live-cell imaging using the same analysis pipeline as for the TetO-LacO dual-array cell line (see description below).

### piggyBac insertion site mapping

The integration sites of the random integrations by PiggyBase were mapped as described in Redolfi et al. ^40^. In short, genomic DNA (2 µg) was fragmented to an average of 500 bp by sonication (Covaris) and ligation of full-length barcoded Illumina adapters was performed using the TruSeq DNA PCR-free kit (Illumina) according to the manufacturer’s guidelines, with the exception that large DNA fragments were not removed. Libraries were pooled together and capture of desired fragments was performed using biotinylated probes against the piggyBac inverted terminal repeats (ITRs) sequences using xGEN Hybridisation reagents (IDT). Following capture, libraries were amplified for 14 cycles (KAPA HiFi Hotstart). Sequencing was performed on the NextSeq500 platform (Illumina) as paired-end 300cycles.

### Capture-Hi-C sample preparation

Capture-Hi-C sample preparation was performed as described previously^3^. In short, for RAD21 depletion, 2×10^7^ cells were treated with 500 nM dTag-13 (Sigma-Aldrich, SML2601-1MG) for 2h at 37°C. All cells were then crosslinked with 1% formaldehyde (EMS, 15710) for 10 min at RT. The reaction was quenched with glycine (final concentration 0.125 M). Lysis was performed in 1 M Tris-HCl pH 8.0, 5 M NaCl and 10% NP40 (Sigma-Aldrich, I8896-50ML) and Complete protease inhibitor (Sigma-Aldrich, 11836170001). Cells were digested using 100 U of MboI (NEB, R0147) and ligated at 16 °C with 10,000 U of T4 DNA ligase (NEB, M0202) in ligase buffer supplemented with 0.8% Triton X-100 (Sigma-Aldrich, T8787) and 240 µg of BSA (NEB, B9000). De-crosslinking was achieved with 400 μg Proteinase K (Macherey Nagel, 740506) at 65°C. The 3C sample was purified using a phenol/chloroform extraction. 3C library preparation and target enrichment using a custom-designed collection of 6979 biotinylated RNA “baits” targeting single MboI restriction fragments chr15:10283500-13195800 (mm9) (**Suppl. Table S2**; Agilent Technologies; as in Ref.^3^) were performed following the SureSelectXT Target Enrichment System for Illumina Paired-End Multiplexed Sequencing Library protocol. However, 9µg of 3C input material instead of 3µg was used for the capture and DNA was sheared using Covaris sonication with the following settings: Duty Factor: 10%; Peak Incident Power (PIP): 175; Cycles per Burst: 200; Treatment Time: 480 seconds; Bath Temperature: 4° to 8°C).

### Targeted nanopore sequencing with Cas9-guided adapter ligation (nCATS)

nCATS was performed as described previously in Zuin et al.^3^ In short, 3-5 gRNAs sequences each (targeting the upstream and downstream regions 2-3 kb external of the respective integration cassette, either LacO or TetO integration) were designed using the IDT online tool https://eu.idtdna.com/site/order/designtool/index/CRISPR_SEQUENCE **(Suppl. Table S3).** Custom designed Alt-R CRISPR-Cas9 crRNAs (3-5 crRNAs targeting the region 5’ and 3-5 crRNAs targeting the region 3’ of the integrated transgene), Alt-R CRISPR-Cas9 tracrRNA (IDT, 1072532) and Alt-R S.p. Cas9 enzyme (IDT, 1081060) were purchased from IDT. Genomic DNA from clones 2G5 and 1F11 was extracted with Gentra Puregene Cell Kit (Qiagen, 158745) following the manufacturer’s instructions. Quality of the High Molecular Weight (HMW) DNA was checked with the TapeStation (Agilent) and 5 µg of HMW DNA were de-phosphorylated using Shrimp Alkaline Phosphatase (rSAP; NEB, M0371) for 30 min at 37°C followed by 5 min at 65°C. To assemble the Alt-R guide RNA duplex (crRNA:tracrRNA), the six Alt-R CRISPR-Cas9 crRNAs were pooled to a final concentration of 100 µM and subsequently incubated in a ratio of 1:1 with 100 µM of Alt-R CRISPR-Cas9 tracrRNA at 95°C for 5 min. 4 pmol of Alt-R S.p Cas9 enzyme were incubated with 8 pmol Alt-R guide RNA (crRNA:tracrRNA) at RT for 20 min to assemble the RNP complex. *In vitro* digestion and A-tailing of the DNA were performed by adding 10 µl of the RNP complex, 10 mM of dATP (NEB, N0440) and 5 U of Taq Polymerase (NEB, M0267) and incubating the samples at 30 min, 37°C followed by 5 min, 72°C. Nanopore sequencing adaptors were ligated using the Ligation Sequencing Kit (Oxford Nanopore Technologies, SQK-CAS109) according to the manufacturer’s instructions. After purification with AMPure PB beads (Beckman Coulter, A63881), samples were loaded into MinION selecting SQL-CAS109 protocol (Oxford Nanopore Technologies).

### Hi-C sample preparation

Hi-C sample preparation was performed as described previously in Redolfi et al.^40^ . Briefly, 6×10^6^ cells were treated with 500 µM auxin (Sigma-Aldrich, I5148-2G) for 90 min and cells were crosslinked with 1% formaldehyde (EMS, 15710) and quenched with 0.125 M glycine for 5min at RT. Cells were lysed in 10 nM Tris-HCl pH 8.0, 10 nM NaCl, 0.2% NP-40 (Sigma-Aldrich, I8896-50ML), complete protease inhibitor (Sigma-Aldrich, 11836170001) and nuclei were digested with 400 U of MboI (NEB, R0147) at 37 °C overnight. End-repair was performed using 40 µM Biotin-11-dATP (Life Technologies, 19524-016) and 50 U DNA Polymerase I Large Klenow fragment (NEB, M0210M) incubating at 37 °C for 45 min. The end-repaired samples were ligated using 10,000 U T4 DNA ligase (NEB, M0202M) in ligase buffer supplemented with 0.8% Triton X-100 (Sigma-Aldrich, T8787) and 120 µg BSA (NEB, B9000) at 16 °C overnight. De-crosslinking was performed by adding 20 µl Proteinase K (20 mg/ml, Macherey-Nagel, 740506) to the ligation mix (1.2 ml) and incubating at 65 °C overnight. DNA was purified using phenol/chloroform and 2 µg of purified 3C sample was sonicated using the Bioruptor Pico (Diagenode). Biotinylated DNA was captured using MyOne Streptavidin T1 magnetic beads (Life Technologies, No. 65601) followed by A-tailing. Library preparation was performed according to NEBNext Ultra DNA Library prep kit instruction (NEB, E7370L) and samples were purified with magnetic AMPure bead (Beckman Coulter, A63881). Hi-C libraries were sequenced on an Illumina Nextseq500 platform (2x42 bp paired-end).

### 4C-seq sample preparation

Sample preparation for 4C-seq was performed as previously described^44^. In short, 10^7^ cells were treated with 500 µM auxin (Sigma-Aldrich, I5148-2G) for 90 min and cross-linked in 2% paraformaldehyde (EMS, 15710) for 10 min and quenched with 0.125 M glycine (final concentration). Lysis was performed in 150 mM NaCl, 50 mM Tris-HCl (pH 7.5), 5 mM EDTA, 0.5% NP-40 (Sigma-Aldrich, I8896-50ML), 1% Triton X-100 (Sigma-Aldrich, T8787). The samples were digested with 200 U DpnII (NEB, R0543M) and subsequently ligated at 16 °C with 50 U T4 DNA ligase (Roche, #10799009001) in a final reaction volume of 7 ml. De- crosslinking was performed with Proteinase K (0.05 μg/µl) at 65 °C and samples were then purified using phenol/chloroform extraction. The second digest was performed with 50 U Csp6I (Thermo Fisher Scientific, ER0211). Samples were ligated with 100 U T4 DNA ligase in a final volume of 14 ml and purified by precipitation in 100% ethanol. The resulting products were used directly as a PCR template for the TetO 4C viewpoint. Primers for PCR were designed according to the set-up used in Redolfi et al^40^ and can be found in **Suppl. Table S3**. Library preparation was performed with the NEBNext Ultra DNA Library prep kit according to the manufacturer’s instructions (NEB, E7370L). 4C-seq libraries were sequenced on an Illumina Hiseq2500 platform (50 cycles, single-end reads).

### Cell cycle stage analysis of RAD21 depleted cells by flow cytometry

To validate that in auxin or dTAG-13 treated cells the distribution of cells in different stages of the cell cycle is not skewed towards cells in S and G2 phase (as cells are arrested in mitosis upon depletion of RAD21), cell cycle stage analysis was performed by flow cytometry. For this, cells were cultured until confluency on a 6-well and treated with the corresponding compound (500 µM auxin (Sigma-Aldrich, I5148-2G) or 500 nM dTAG-13 (Sigma-Aldrich, SML2601-1MG) resuspended in culturing medium) for the time indicated (0h, 1.5h, 2h, 6h). The cells were then harvested and 3×10^6^ cells were fixed in 4% paraformaldehyde (EMS, 15710) at RT for 15 min. The cells were washed in PBS and stained with 5 µg/ml 4′,6-Diamidino-2- phenylindole (DAPI) (D9564-10MG, Sigma-Aldrich) in 1xPBS+0.1% Triton X-100 (Sigma-Aldrich, T8787) for 30 min at RT. The cells were analyzed with a 405 nm laser line at BD LSR II SORP Analyser (BD Biosciences). Distributions of cell cycle stage profiles were analyzed using FlowJo (v10, BD Biosciences).

### Western Blot

To validate the targeted degradation of RAD21, CTCF and WAPL in the degron cell lines, cells were cultured on a 6-well to confluency and degradation was induced by adding 500 nM dTag-13 (Sigma-Aldrich, SML2601-1MG) or 500 µM auxin (Sigma-Aldrich, I5148-2G) and incubating for the indicated time (0h, 1.5h, 2h, 6h, 24h) at 37°C, 8% CO2. The cells were washed twice with PBS and lysed in 200µl RIPA buffer (150 mM sodium chloride, 1.0% NP-40 (Sigma-Aldrich, I8896-50ML), 0.5% sodium deoxycholate, 0.1% sodium dodecyl sulfate (SDS), 50 mM Tris pH 8.0) incubating 10 min at 4°C. Lysed cells were frozen in liquid nitrogen. For the SDS-PAGE, cell lysates were thawed on ice and 2.5 U/ml SuperNuclease (Sino Biological Inc, SSNP01) was added to digest DNA for 10 min. Protein levels were quantified using the Pierce BCA protein assay (Thermo Fisher Scientific, 23225). 10 µg of protein extract were loaded onto a Mini-PROTEAN TGX Precast Gel (4-15% gradient, BioRad, 4561086) in 1xLaemmli buffer (BioRad, 1610747) containing 100 mM DTT. Samples were run at 180 V for 90 min in Tris-Glycine buffer (25 mM Tris pH 8.3, 192 mM Glycine, 0.1% (w/v) SDS). The protein was transferred to a Trans-Blot Turbo Mini 0.2 µm Nitrocellulose membrane (BioRad, 1704158) using the Trans-Blot Turbo Transfer System from BioRad. The membrane was blocked for 1h at RT (shaking) in Odyssey Blocking Buffer (PBS) (Li-Cor Biosciences, 927-40000). The membrane was incubated with 1 µg/ml primary antibody (rabbit polyclonal anti-RAD21, abcam ab154769, rabbit polyclonal anti-WAPL, Proteintech 16370-1-AP, rabbit polyclonal anti-CTCF, Cell Signaling Technologies #2899S, rabbit anti-PK-tag (V5), abcam ab15828, and mouse monoclonal anti-Tubulin, (DM1A), Cell Signaling Technologies #3872) in Odyssey Blocking Buffer (PBS) overnight at 4°C. The membrane was washed three times in PBS+0.1%Tween-20 for 5min each shaking and then incubated for 1h at RT (shaking) with secondary antibodies (IRDye 800CW Goat anti-rabbit IgG and IRDye 680RD Goat anti-Mouse IgG, dilution 1:10,000, Li-Cor BioSciences 926-32211 and 926-68070) in Odyssey Blocking Buffer (PBS). The membrane was washed three times in PBS+0.1%Tween-20 and imaged on the Odyssey infrared imaging system (Li-Cor Biosciences).

### Live-cell imaging

35mm glass-bottom dishes (Mattek, P35G-1.5-14-C) were coated with 1-2µg/ml Laminin (Sigma-Aldrich, L2020) in PBS at 37°C overnight. Cells (1×10^6^) were seeded in Fluorobrite medium (as described above) 24h before imaging. For targeted degradation of RAD21, WAPL or CTCF in the degron cell lines, the medium was exchanged to medium containing 500 µM auxin (Sigma-Aldrich, I5148-2G) at the respective time required for complete degradation of the protein target prior to imaging (RAD21: 90 min, WAPL: 24h, CTCF: 6h). For targeted depletion of RAD21 using the FKBP degron system (dual array cell lines), cells were cultured in Fluorobrite medium containing 500 nM dTAG-13 (Sigma-Aldrich, SML2601-1MG) 2h prior to imaging.

For fixed cell measurements to estimate the localization error, 1×10^6^ cells were seeded onto Mattek dishes and incubated for 24h at 37°C, 8%CO2. The medium was removed and the cells were fixed in 4% paraformaldehyde (Electron Microscopy Sciences, 15710) in PBS for 10 min at RT. The cells were washed three times in PBS and Fluorobrite medium was added to the Mattek dish to achieve comparable background fluorescence levels.

Cells were imaged with a Nikon Eclipse Ti-E inverted widefield microscope equipped with a Total Internal Reflection Microscopy iLAS2 module (Roper Scientific), a Perfect Focus System (Nikon) and motorized Z-Piezo stage (ASI) using a CFI APO TIRF 100 x 1.49 NA oil immersion objective (Nikon). The microscope was operating in highly inclined and laminated optical sheet (HILO) mode^26^. Excitation sources were a 488 nm, 200 mW Toptica iBEAM SMART laser and a 561 nm 200 mW Coherent Sapphire laser. Images were collected on two precisely aligned back-illuminated Evolve 512 Delta EMCCD cameras with a pixel size of 16 μm x 16 μm (Photometrics). Cells were maintained at 37 °C and 8% CO2 using an enclosed microscope environmental control setup (The BOX and The CUBE, Life Science Instruments). Before the acquisition of movies for the dual-array set-up, TetraSpeck™ Microspheres, 0.1 µm beads (Thermo Fisher Scientific, T7279) were imaged to allow for correction of chromatic aberrations during image processing and analysis. Movies for measurement of random TetO integrations in degron cell lines were acquired every 10 s (exposure time: 50 ms) in 34 z-planes (10 µm stack, dz = 300 nm) with the Visiview software (Visiview 4.4.0.12, Visitron). Images for measurement of cell lines with the dual array set-up were acquired every 30 s, with an exposure time of 50 ms, respectively, each in a sequential mode with 21 z-planes (6 µm stack, dz = 300 nm). For the measurement of the time it takes the operator arrays to displace by its own size, images were acquired continuously on a single focal plane over 10 s every 0.1 s with exposure times of 50 ms.

### Image processing

Raw images were deconvolved using the Huygens Remote Manager and a classical maximum likelihood estimation algorithm with a theoretical point-spread-function. The initial signal-to-noise ratios were estimated from the images and images were deconvolved until one of the following stopping criteria was reached: The maximum number of iterations were performed (for random integrations: 20 cycles, for tdTomato and eGFP; in dual-color set-up: 15 cycles for tdTomato signal, for GFP signal: 5 cycles) or a quality change criterion below 0.001 was returned. Representative image series shown in the main figures were deconvolved as described above, adjusted to display the same brightness and contrast and interpolated using a bicubic interpolation. Movies were corrected for bleaching over time using an exponential fit. The 2D projection of intensity changes over time was created using the the Temporal Color Code in Fiji (https://github.com/fiji/fiji/blob/master/plugins/Scripts/Image/Hyperstacks/Temporal-Color_Code.ijm).

### Multi operator image analysis and localisation error estimation

We used deepBlink^27^, a convolutional neural network-based spot detection algorithm, for spot detection. We employed custom models trained on a combination of the following datasets: smFISH and SunTag datasets provided by deepBlink and in-house manually curated live cell imaging images. The models can be found https://github.com/zhanyinx/SPT_analysis/tree/main/models. The parameters and models used for each cell line can be found in **Suppl. Table S4.** To detect 3D spots, we applied deepBlink to all z-stacks separately followed by linkage of the spots across z-stacks using Trackpy^45^; the precise 3D coordinates of the spots are then determined using 3D gaussian fitting using a voxel of size 6x6x4 pixels centered at the spot in the brightest z stack. 3D spots are fed into TrackMate for tracking using linear assignment problem (LAP) tracker. Each track is assigned to manually annotated cell masks (from max z-projection of frame 93) using a custom script (https://github.com/zhanyinx/SPT_analysis/blob/main/source/spot_detection_tracking/assign_cellids.py), which uses the majority rule. Motion correction is then performed using a roto-translation model. Briefly, for each pair of consecutive frames, matches spots (i.e. same spot across time) are determined by solving the linear assignment problem using scipy.optimize.linear_sum_assignment function with the euclidean distance between spots as measure of distance. The roto-translation model is calculated for each pair of consecutive frames using the matched spots.

Tracks with less than 10 spots are filtered out for followed up analysis. To calculate the mean square displacement (MSD), we first calculate the time-averaged MSD for each trajectory. We then calculate the ensemble average (across trajectories) MSD by pooling all replicates. We corrected the localisation error effect on the MSD curve by estimating the standard deviation of the error distribution using fixed images as described in Ref.^46^. To calculate the scaling (*α*) and the generalized diffusion coefficient (D) of each MSD curve, we fitted the ensemble average of the log-time average MSD between 10-100s. To test the significance of differences between conditions, we fitted *α* and diffusion coefficient for each cell. The p-value is calculated using Student t-test (two-sided). Since we are always comparing two conditions whose cell cycle profiles are similar, we ignore the effect of sister chromatids. All scripts used for the analysis can be found at https://github.com/zhanyinx/SPT_analysis/. The specific Fiji and relative plug-ins can be found at https://github.com/giorgettilab/Mach_et_al_chromosome_dynamics/tree/master/Fiji.

### Dual color image analysis and radial localisation uncertainty

Spot detection and tracking is done as for the multi operator image analysis. We correct for chromatic aberration by computing the roto-translation on beads taken the same exact day of sample imaging. Briefly, beads spots are detected using deepBlink and matched across channels by solving the linear assignment problem using scipy.optimize.linear_sum_assignment function with the euclidean distance between spots as measure of distance. We used the set of matched beads spots across channels to compute the roto-translation. We apply the roto-translation to the sample images to correct for chromatic aberration. To increase the ability to detect longer tracks, we used an in-house script to stitch multiple tracks belonging to the same cell (https://github.com/zhanyinx/SPT_analysis/blob/main/source/dual_channel_analysis/utils.py, stitch function). In short, if two tracks overlap more than 50% in time, the shortest one is filtered out. We called cell masks using CellPose^47^ on the max z-projection of the middle frame of the movie using GFP channel. For tracks with time-overlaps lower than 50%, the overlapping part of the tracks are randomly removed from one of the two tracks. The resulting tracks are stitched if the distance across the time gap is smaller than 1.6µm. To match tracks across channels, we used the following measure to calculate the distance between tracks across channels

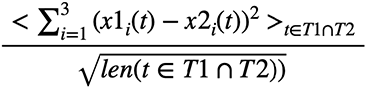

Where x1 are the coordinates from channel1 and x2 are the coordinates from channel2, T1 contains all the time frames from channel1 and T2 contains all the time frames from channel2, and len is a function that returns the length of an array. We solved the linear assignment problem using the distance measure above to match tracks across channels. Tracks with average distances across channels higher than 1µm are filtered out. Matched tracks with lower than 25 time points are filtered out. For each matched pair of tracks, we calculate the pairwise distance using the euclidean distance in 3D. We define noisy pairwise distance using the ratio of the pairwise distance in 3D and 2D. In particular, we defined noisy the top 5% of this ratio and filtered them out. To calculate the radial mean square displacement (MSD), we first calculate the time-averaged radial MSD for each pairwise distance “trajectory”. We then calculate the ensemble average (across trajectories) of the log of time-averaged radial MSD. We corrected for the radial localisation uncertainty by estimating the standard deviation of the error distribution using fixed images as described in Ref.^46^. To calculate the scaling (*α*) and the generalized diffusion coefficient (D) of each MSD curve, we fitted the ensemble average time average MSD between 30-300s. Since we are always comparing two conditions whose cell cycle profiles are similar, we ignore the effect of sister chromatids. All scripts used for the analysis can be found at https://github.com/zhanyinx/SPT_analysis/.

### Estimation of experimental uncertainty on radial distance

To estimate our uncertainty in detecting distances across channels, we used a cell line with multiple integration of TetO arrays that can be tagged with TetR-eGFP and TetR-tdTomato. Spot detection is done as for our dual color lines. We corrected for chromatic aberration using TetraSpeck™ Microspheres, 0.1 µm beads (Thermo Fisher Scientific, T7279) and then matched spots across channels by solving the linear assignment problem using scipy.optimize.linear_sum_assignment function with the euclidean distance between spots as measure of distance. Spots across channels with distances higher than a threshold are filtered out. We used a threshold of 300 nm for matching the spots registration. We applied a second round of chromatic aberration correction using the set of registered points itself. The resolution limit (uncertainty) is then estimated as the average distance between registered spots which corresponds to 130 nm +/- 70 nm.

### Estimation of sister chromatids

To estimate the probability of encountering sister chromatids in our dual color experiments, we manually labeled around 1400 images randomly sampled from all movies. We assigned spots to cell masks detected using CellPose. We estimated that sister chromatids occur in ∼3% of the cells.

### Hidden markov model for detection of proximal state

To detect the proximal state in a threshold independent manner, we used a hidden markov model (HMM) with two hidden states (proximal and distal). We used a Gaussian model for the emission probabilities. Only distance trajectories with less than 20% missing values at any time point are kept. Missing values are filled with the first preceding time point with distance value. In order to more reliably detect the proximal state, we used all the trajectories from the experimental condition with both cohesin and CTCF to train an HMM. We then re-train an HMM model per experimental condition by using the proximal state (gaussian mean and standard deviation) from the experimental condition with both cohesin and CTCF. Finally we applied the experimental condition specific-HMM to every trajectory to estimate the contact duration and rate of contact formation for all the experimental conditions. The HMM model training can be found as a jupyter notebook (https://github.com/zhanyinx/SPT_analysis/blob/main/notebooks/HMM_experimental_data.ipynb). We modified the hmmlearn library to allow fixing proximal state during HMM training. The modified hmmlearn library can be found at https://github.com/zhanyinx/hmmlearn

### Median convergent CTCF distance

To calculate the average distance between convergent CTCF within TADs, we took the list of CTCF motifs from Nora et al 2017^9^. The list of TADs has been taken from Zuin, Roth et al 2022^3^. By keeping only pairs of convergent CTCF motifs within TADs, the average distance is estimated to be 140932 bp.

### Analysis of PiggyBac integration mapping

To exclude reads coming from the TetR or LacI, the two ends of paired end reads were mapped separately to the piggyBac-TetR and LacI sequences (https://github.com/giorgettilab/Mach_et_al_chromosome_dynamics/tree/master/sequences) using QuasR (qAlign). Only unmapped reads were kept and mapped to the piggybac-TetO array sequence (https://github.com/giorgettilab/Mach_et_al_chromosome_dynamics/tree/master/sequences). Hybrid pairs with one of the read-end mapping to array were kept. The second reads from hybrid pairs were mapped to the mouse genome (build mm9) using QuasR (qAlign). Reads were then piled up in 25-bp windows using csaw (windowCounts function). Integration sites can be identified because they correspond to local high-read coverage. Local coverage was calculated by resizing all non-zero 25-bp windows up to 525 bp (expanding by 250 bp upstream and downstream). Overlapping windows were then merged using reduce (from GenomicRanges), resulting in a set of windows {wi}. The size distribution of wi is multimodal, and only wi from the second mode onward were kept. For each wi we estimated the coverage ci as the number of non-zero 25-bp windows. Only {wi} where the coverage was >16 were considered. The exact positions of the integration sites were then identified with the center of wi.

### Hi-C analysis

Hi-C data were analyzed using HiC-Pro version 3.1.0^48^ with the --very-sensitive --end-to-end --reorder options. Only unique reads were mapped to the mouse genome (build mm9). Contact maps were combined at 8 kb with iterative correction applied afterwards. (https://github.com/giorgettilab/Mach_et_al_chromosome_dynamics/tree/master/Hi-C)

### Loop analysis

Loops were called on Hi-C map of WT sample using Mustache^49^ version 1.0.1 on 8 kb contact map, with p-value threshold 0.1 and sparsity threshold 0.75. Pileup analysis was performed for all samples based on WT loops using coolpup.py version 0.9.2 ^50^

### 4-C analysis

First, we trimmed the connector sequence, then we filtered out overrepresented sequences. We mapped sequences to the mouse mm9 genome using a custom R script (https://github.com/giorgettilab/Mach_et_al_chromosome_dynamics/).

### Nanopore sequencing analysis

To map nanopore sequencing reads, we first built a custom ‘genome’ consisting of the LacO or TetO cassette flanked by ∼2.5kb mouse genomic sequence upstream and downstream of the target integration site. The custom genome can be found at (https://github.com/giorgettilab/Mach_et_al_chromosome_dynamics/tree/master/nanopore). Analysis has been performed as in Zuin et al.^3^. Briefly, reads were mapped to the custom genome using minimap2 (v. 2.17-r941) with “-x map-ont” parameter. Nanopore sequencing analysis has been implemented using Snakemake workflow (v. 3.13.3). Reads were visualized using IGV (v. 2.9.4). Full workflow can be found at https://github.com/zhanyinx/Zuin_Roth_2021/tree/main/Nanopore.

### Capture-C analysis

capture-C data were processed as in Zuin et al.^3^ using HiC-Pro^48^ (v. 2.11.4). Briefly, read pairs were mapped to the mouse genome (build mm9). Chimeric reads were recovered after recognition of the ligation site. Only unique valid pairs mapping to the target regions were used to build contact maps. Iterative correction (ICE)^51^ was then applied on binned data. The target regions can be found at https://github.com/zhanyinx/Zuin_Roth_2021

### Differential Capture-C maps

We accounted for differences in genomic distances due to the presence of the ectopic sequence when evaluating the structural perturbation induced by the insertion of the TeTO and LacO arrays. To account for these differences, we generated distance-normalized capture-C maps where each entry corresponds to the interaction normalized by the corrected genomic distance between the interacting bins. We then calculated the WT-allele corrected ratios between distance normalized maps using the following formula:

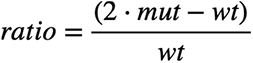

Where *mut* is the distance normalized heatmap of the line with TeTO or LacO arrays and *wt* is the distance normalized heatmap of the wild-type. A bilinear smoothing with a window of 2 bins has been applied to the ratio maps to evaluate the structural perturbation induced by the insertion of the arrays and the effect of ectopic CTCF sites.

### Simulations

Polymer simulations were performed using LAMMPS^52^. We chose Langevin dynamics with the NVT thermostat. Arbitrary units were set such that thermal energy *k_B_T* = 1, where k_B -– the Boltzmann constant and T – temperature, corresponding to 300K. To simulate the loop extrusion process, we developed and embedded in LAMMPS a new package called “USER-LE”. Loop extrusion model contains extruders and barriers on the polymer. An extruder is represented as an additional sliding bond, which extrudes the loop in a two-sided manner. It can be loaded to the polymer between (i) and (i+2) beads with a certain probability only when the bead (i+1) is unoccupied by another extruder and is not a barrier. Each extruder can be unloaded from polymer with a certain probability. Every bead could be occupied by only one extruder. Extruders cannot pass through each other. When extruders meet each other on the polymer, they stall until one of them is released. Every extruder attempts to make an extruding step every N simulation steps.

In addition to “neutral” polymer beads, there are 3 types of barriers blocking loops coming from the left, from the right and from any direction. These barriers mimic CTCF sites, for which one can define a probability for the loop extruder to go through (the same probability for all barriers). To launch loop extrusion, one should define three fixes with LAMMPS syntax: loading, unloading and loop extrusion. Loading: frequency in number of steps to try to load extruders, types of beads, max distance to create, type of the bond (extruder) to be created, probability to create, seed for pseudorandom generator of numbers, new type of the first beads and new type for the second bead. Unloading: frequency in number of steps to try to unload extruders, type of the bond (extruder), min distance to release bond, probability to release bond, seed for pseudorandom number generator. Loop extrusion: frequency in number of steps to try to move extruders, neutral polymer type, left barrier type, right barrier type, probability to go through the barrier, type of the bond (extruder), and type of two-sided barrier (optional).

### Conversion of simulation timestep to real time

In order to convert the timescales between experiments and simulations, we measured the time needed for a bead (and 8 kb TetO array) to move for its own size (95 nm = 15 nm x = estimated size of a nucleosome x estimated number of nucleosomes in each 8kb segment, assuming that chromatin inside each 8kb behave as an ideal chain; this is in line with the previous estimations for a 3 kb segment^16^). To this aim, 2D movies acquired in cells with randomly inserted TetOs every 0.1 second for 10s were analyzed with the same strategy as in **Multi-operator image analysis and localisation error estimation**, now without 3D linkage. We used the MSD curve to estimate the time necessary to move for 95 nm for each line. We used the average value across 3 lines to convert the timescales between experiments and simulations.

In simulations, a bead needs approx. 1800 simulation steps to move of its own size, whereas in experiment this time is estimated to be approx. 0.86 sec. Therefore 1 sec of the real time corresponds to approx. 2100 simulation steps.

### Metrics for quantifying similarity between simulations and experiment

To evaluate the similarity between experimental features (f) (here: contact duration, probability of being in the proximal state, fraction of time spent in the proximal state) we used the relative difference defined as:

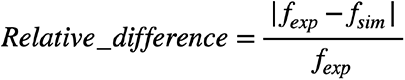

To find the set of loop extrusion parameters that best reproduce the experiments in the presence of cohesin, we used the sum of the relative difference of the following features: gaussian mean of contact state, the contact duration and the fraction of time spent in the looped state in the +/- CTCF. To find the set of loop extrusion parameters that best reproduce the experiments in the absence of cohesin, we decreased the loading rate to reach extruder densities of 0.87 per Mb while keeping the extruder residence time constant (5.5 min).

### Data availability

All capture-C, Hi-C, 4C, integration site mapping sequencing fastq files generated in this study have been uploaded to the Gene Expression Omnibus (GEO) under accession GSE197238. The following public databases were used: BSgenome.Mmusculus.UCSC.mm9 (https://bioconductor.org/packages/release/data/annotation/html/BSgenome.Mmusculus.UCSC.mm9.html).

### Code availability

Custom codes generated in this study are available at: https://github.com/zhanyinx/SPT_analysis/ (image analysis); https://github.com/giorgettilab/Mach_et_al_chromosome_dynamics/ (4C,Hi-C, nanopore, simulation analysis); the modified version of hmmlearn can be found at https://github.com/zhanyinx/hmmlearn.

## REFERENCES

1. Marchal, C., Sima, J. & Gilbert, D. M. Control of DNA replication timing in the 3D genome. Nat. Rev. Mol. Cell Biol. 20, 721–737 (2019).

2. Arnould, C. et al. Loop extrusion as a mechanism for formation of DNA damage repair foci. Nature 590, 660–665 (2021).

3. 3. Zuin, J. et al. Nonlinear control of transcription through enhancer-promoter interactions. bioRxiv 2021.04.22.440891 (2021) doi:10.1101/2021.04.22.440891.

4. Nora, E. P. et al. Spatial partitioning of the regulatory landscape of the X-inactivation centre. Nature 485, 381–385 (2012).

5. Dixon, J. R. et al. Topological domains in mammalian genomes identified by analysis of chromatin interactions. Nature 485, 376–380 (2012).

6. Fudenberg, G. et al. Formation of Chromosomal Domains by Loop Extrusion. Cell Rep. 15, 2038–2049 (2016).

7. Nora, E. P. et al. Molecular basis of CTCF binding polarity in genome folding. Nat. Commun. 11, 5612 (2020).

8. Li, Y. et al. The structural basis for cohesin–CTCF-anchored loops. Nature 578, 472–476 (2020).

9. Nora, E. P. et al. Targeted Degradation of CTCF Decouples Local Insulation of Chromosome Domains from Genomic Compartmentalization. Cell 169, 930–944.e22 (2017).

10. Wutz, G. et al. Topologically associating domains and chromatin loops depend on cohesin and are regulated by CTCF, WAPL, and PDS5 proteins. EMBO J. 36, 3573–3599 (2017).

11. Rao, S. S. P. et al. Cohesin Loss Eliminates All Loop Domains. Cell 171, 305–320.e24 (2017).

12. Haarhuis, J. H. I. et al. The Cohesin Release Factor WAPL Restricts Chromatin Loop Extension. Cell 169, 693–707.e14 (2017).

13. McCord, R. P., Kaplan, N. & Giorgetti, L. Chromosome Conformation Capture and Beyond: Toward an Integrative View of Chromosome Structure and Function. Mol. Cell 77, 688–708 (2020).

14. Hansen, A. S., Cattoglio, C., Darzacq, X. & Tjian, R. Recent evidence that TADs and chromatin loops are dynamic structures. Nucleus 9, 20–32 (2018).

15. Schoenfelder, S. & Fraser, P. Long-range enhancer–promoter contacts in gene expression control. Nat. Rev. Genet. 20, 437–455 (2019).

16. Giorgetti, L. et al. Predictive Polymer Modeling Reveals Coupled Fluctuations in Chromosome Conformation and Transcription. Cell 157, 950–963 (2014).

17. Bintu, B. et al. Super-resolution chromatin tracing reveals domains and cooperative interactions in single cells. Science 362, eaau1783 (2018).

18. Finn, E. H. & Misteli, T. Molecular basis and biological function of variability in spatial genome organization. Science 365, eaaw9498 (2019).

19. Davidson, I. F. et al. DNA loop extrusion by human cohesin. Science 366, 1338–1345 (2019)

20. Hansen, A. S., Pustova, I., Cattoglio, C., Tjian, R. & Darzacq, X. CTCF and cohesin regulate chromatin loop stability with distinct dynamics. eLife https://elifesciences.org/articles/25776(2017) doi:10.7554/eLife.25776.

21. Gabriele, M. et al. Dynamics of CTCF and cohesin mediated chromatin looping revealed by live-cell imaging. 2021.12.12.472242 https://www.biorxiv.org/content/10.1101/2021.12.12.472242v1(2021) doi:10.1101/2021.12.12.472242.

22. Cadiñanos, J. & Bradley, A. Generation of an inducible and optimized piggyBac transposon system†. Nucleic Acids Res. 35, e87 (2007).

23. Dubarry, M., Loïodice, I., Chen, C. L., Thermes, C. & Taddei, A. Tight protein–DNA interactions favor gene silencing. Genes Dev. 25, 1365–1370 (2011).

24. Krebs, A. R. et al. Genome-wide Single-Molecule Footprinting Reveals High RNA Polymerase II Turnover at Paused Promoters. Mol. Cell 67, 411–422.e4 (2017).

25. Liu, N. Q. et al. WAPL maintains a cohesin loading cycle to preserve cell-type-specific distal gene regulation. Nat. Genet. 53, 100–109 (2021).

26. Tokunaga, M., Imamoto, N. & Sakata-Sogawa, K. Highly inclined thin illumination enables clear single- molecule imaging in cells. Nat. Methods 5, 159–161 (2008).

27. Eichenberger, B. T., Zhan, Y., Rempfler, M., Giorgetti, L. & Chao, J. A. deepBlink: threshold- independent detection and localization of diffraction-limited spots. Nucleic Acids Res. 6, 7292–7297 (2021)

28. Gu, B. et al. Transcription-coupled changes in nuclear mobility of mammalian cis-regulatory elements. Science 359, 1050–1055 (2018).

29. Khanna, N., Zhang, Y., Lucas, J. S., Dudko, O. K. & Murre, C. Chromosome dynamics near the sol-gel phase transition dictate the timing of remote genomic interactions. Nat. Commun. 10, 2771 (2019).

30. Cattoglio, C. et al. Determining cellular CTCF and cohesin abundances to constrain 3D genome models. eLife 8, e40164 (2019).

31. Holzmann, J. et al. Absolute quantification of cohesin, CTCF and their regulators in human cells. eLife 8, e46269 (2019).

32. Tamm, M. V. & Polovnikov, K. Dynamics of polymers: classic results and recent developments. ArXiv170709885 Cond-Mat Q-Bio (2017).

33. Jung, I. et al. A Compendium of Promoter-Centered Long-Range Chromatin Interactions in the Human Genome. Nat. Genet. 51, 1442–1449 (2019).

34. Gilpatrick, T. et al. Targeted nanopore sequencing with Cas9-guided adapter ligation. Nat. Biotechnol. 38, 433–438 (2020).

35. Nabet, B. et al. The dTAG system for immediate and target-specific protein degradation. Nat. Chem. Biol. 14, 431–441 (2018).

36. Wutz, G. et al. ESCO1 and CTCF enable formation of long chromatin loops by protecting cohesinSTAG1 from WAPL. eLife 9, e52091 (2020).

37. Szabo, Q. et al. Regulation of single-cell genome organization into TADs and chromatin nanodomains. Nat. Genet. 52, 1151–1157 (2020).

38. Liu, N. Q. et al. Rapid depletion of CTCF and cohesin proteins reveals dynamic features of chromosome architecture. 2021.08.27.457977 https://www.biorxiv.org/content/10.1101/2021.08.27.457977v1(2021) doi:10.1101/2021.08.27.457977.

39. Masui, O. et al. Live-Cell Chromosome Dynamics and Outcome of X Chromosome Pairing Events during ES Cell Differentiation. Cell 145, 447–458 (2011).

40. Redolfi, J. et al. DamC reveals principles of chromatin folding in vivo without crosslinking and ligation. Nat. Struct. Mol. Biol. 26, 471–480 (2019).

41. Pollex, T. & Heard, E. Nuclear positioning and pairing of X-chromosome inactivation centers are not primary determinants during initiation of random X-inactivation. Nat. Genet. 51, 285–295 (2019).

42. Lau, I. F. et al. Spatial and temporal organization of replicating Escherichia coli chromosomes. Mol. Microbiol. 49, 731–743 (2003).

43. Grimm, J. B. et al. A general method to fine-tune fluorophores for live-cell and in vivo imaging. Nat. Methods 14, 987–994 (2017).

44. Splinter, E., de Wit, E., van de Werken, H. J. G., Klous, P. & de Laat, W. Determining long-range chromatin interactions for selected genomic sites using 4C-seq technology: From fixation to computation. Methods 58, 221–230 (2012).

45. Allan, D. B., Caswell, T., Keim, N. C., van der Wel, C. M. & Verweij, R. W. soft-matter/trackpy: Trackpy v0.5.0. (Zenodo, 2021). doi:10.5281/zenodo.4682814.

46. Kepten, E., Bronshtein, I. & Garini, Y. Improved estimation of anomalous diffusion exponents in single- particle tracking experiments. *Phys*. Rev. E 87, 052713 (2013).

47. Stringer, C., Wang, T., Michaelos, M. & Pachitariu, M. Cellpose: a generalist algorithm for cellular segmentation. Nat. Methods 18, 100–106 (2021).

48. Servant, N. et al. HiC-Pro: an optimized and flexible pipeline for Hi-C data processing. Genome Biol. 16, 259 (2015).

49. Roayaei Ardakany, A., Gezer, H. T., Lonardi, S. & Ay, F. Mustache: multi-scale detection of chromatin loops from Hi-C and Micro-C maps using scale-space representation. Genome Biol. 21, 256 (2020).

50. Flyamer, I. M., Illingworth, R. S. & Bickmore, W. A. Coolpup.py: versatile pile-up analysis of Hi-C data. Bioinformatics 36, 2980–2985 (2020).

51. Imakaev, M. et al. Iterative correction of Hi-C data reveals hallmarks of chromosome organization. Nat. Methods 9, 999–1003 (2012).

52. Thompson, A. P. et al. LAMMPS - a flexible simulation tool for particle-based materials modeling at the atomic, meso, and continuum scales. Comput. Phys. Commun. 271, 108171 (2022).

